# Molecular interactions of PCSK9 with an inhibitory nanobody, CAP1 and HLA-C: functional regulation of LDLR levels

**DOI:** 10.1101/2022.10.20.513093

**Authors:** Carole Fruchart Gaillard, Ali Ben Djoudi Ouadda, Lidia Ciccone, Emmanuelle Girard, Sepideh Mikaeeli, Alexandra Evagelidis, Maïlys Le Dévéhat, Delia Susan-Resiga, Evelyne Cassar Lajeunesse, Hervé Nozach, Oscar Henrique Pereira Ramos, Aurélien Thureau, Pierre Legrand, Annik Prat, Vincent Dive, Nabil G. Seidah

## Abstract

**Objective:** The liver-derived circulating PCSK9 enhances the degradation of the LDL receptor (LDLR) in endosomes/lysosomes. PCSK9 inhibition or silencing is presently used in clinics worldwide to reduce LDL-cholesterol, resulting in lower incidence of cardiovascular disease and possibly cancer/metastasis. The mechanism by which the PCSK9-LDLR complex is sorted to degradation compartments is not fully understood. We previously suggested that out of the three M1, M2 and M3 subdomains of the C-terminal Cys/His-rich-domain (CHRD) of PCSK9, only M2 is critical for the activity of extracellular of PCSK9 on cell surface LDLR. This likely implicates the binding of M2 to an unknown membrane-associated “protein X” that would escort the complex to endosomes/lysosomes for degradation. We reported that a nanobody P1.40 binds the M1 and M3 domains of the CHRD and inhibits the function of PCSK9. It was also reported that the cytosolic adenylyl cyclase-associated protein 1 (CAP1) could bind M1 and M3 subdomains and enhance the activity of PCSK9. In this study, we determined the 3-dimensional structure of the CHRD-P1.40 complex to understand the intricate interplay between P1.40, CAP1 and PCSK9 and how they regulate LDLR degradation.

**Methods:** X-ray diffraction of the CHRD-P1.40 complex was analyzed with a 2.2 Å resolution. The affinity and interaction of PCSK9 or CHRD with P1.40 or CAP1 was analyzed by atomic modeling, site-directed mutagenesis, bio-layer interferometry, expression in hepatic cell lines and immunocytochemistry to monitor LDLR degradation. The CHRD-P1.40 interaction was further analyzed by deep mutational scanning and binding assays to validate the role of predicted critical residues. Conformational changes and atomic models were obtained by small angle X-ray scattering (SAXS).

**Results:** We demonstrate that PCSK9 exists in a closed or open conformation and that P1.40 favors the latter by binding key residues in the M1 and M3 subdomains of the CHRD. Our data show that CAP1 is well secreted by hepatic cells and binds extracellular PCSK9 at distinct residues in the M1 and M3 domains and in the acidic prodomain. CAP1 stabilizes the closed conformation of PCSK9 and prevents P1.40 binding. However, CAP1 siRNA only partially inhibited PCSK9 activity on the LDLR. By modeling the previously reported interaction between M2 and an **R**-X-**E** motif in HLA-C, we identified Glu_567_ and Arg_549_ as the critical M2 residues binding HLA-C. Amazingly, these two residues are also required for the PCSK9-induced LDLR degradation.

**Conclusions:** The present study reveals that CAP1 enhances the function of PCSK9, likely by twisting the protein into a closed configuration that exposes the M2 subdomain needed for targeting the PCSK9-LDLR complex to degradation compartments. We hypothesize that “protein X”, which is expected to guide the LDLR-PCSK9-CAP1 complex to these compartments after endocytosis into clathrin-coated vesicles, is HLA-C or a similar MHC-I family member. This conclusion is supported by the PCSK9 natural loss-of-function Q554E and gain-of-function H553R M2 variants, whose consequences are anticipated by our modeling.

## 1. INTRODUCTION

Liver-derived PCSK9 enhances the levels of circulating LDL-cholesterol (LDLc) [1-4] by promoting the degradation of the LDL receptor (LDLR) [5] within the acidic endosomal/lysosomal pathway [6-8]. The crystal structures of PCSK9 revealed the existence of three distinct structural domains playing critical roles in the regulation of PCSK9 and its intracellular traffic: the prodomain (amino acids (aa), 31-152), the catalytic subunit (aa 152-421), and the C-terminal Cys/His-rich domain (CHRD; aa 453-692) [9,10] that is composed of three repeat modules M1 (aa 453-529), M2 (aa 530-603), and M3 (aa 604-692) [9]. Structural [9-12] and biochemical [13] studies demonstrated that the catalytic domain of PCSK9 binds the epidermal growth factor-A (EGF-A) domain of the LDLR, and the PCSK9-LDLR complex is internalized into clathrin-coated vesicles [14-16]. This complex is then directed toward endosomes/lysosomes for degradation by an undefined mechanism(s), preventing LDLR recycling to the cell surface [6,14,15,17]. Since removal of the cytosolic tail of the LDLR or its replacement with that of another protein did not affect the activity of PCSK9 [14,18-22], this suggested that an unknown membrane-bound protein, termed “protein X” [23], may bind the critical M2 domain of the CHRD [24] and escort the PCSK9-LDLR complex to lysosomal degradative compartments [25,26]. P1.40, a CHRD-specific single domain antibody (nanobody), prevents the ability of PCSK9 to enhance the degradation of the LDLR in cells [27] and in transgenic mice that specifically express human PCSK9 [28].

The widely expressed cytosolic adenylyl cyclase-associated protein 1 (CAP1) that acts as a receptor to resistin [29], was recently shown to bind the CHRD and to significantly enhance the degradation of the PCSK9-LDLR complex [30]. However, how the soluble cytosolic CAP1 meets extracellular PCSK9 and enhances the lysosomal degradation of the complex LDLR-PCSK9-CAP1 remains unclear.

Very recently, PCSK9 was shown to induce the lysosomal degradation of the type-I membrane-bound major histocompatibility complex-I (MHC-I), *via* an interaction with the M2 domain of the CHRD, independently from the LDLR [31]. In contrast, PCSK9 interacts with the LDLR primarily through its catalytic domain [9-13].

Herein, we first show that the cytosolic CAP1 is secreted. X-ray structure analysis and modeling identified the distinct residues in the M1 and M3 domains of the CHRD that interact with the inhibitory P1.40. We also defined residues within PCSK9’s prodomain as well as M1 and M3 subdomains involved in CAP1 interaction. Our structural, biochemical, and immunocytochemical data support the notion that “protein X” is likely composed of at least two CHRD-binding components, including CAP1 that maximizes the exposure of the M2 domain to an undefined “protein X” that binds the M2 domain and sorts the PCSK9-LDLR complex to endosomes/lysosomes for degradation. Our mutagenesis and modeling data further suggest that HLA-C is a likely “protein X” candidate in hepatocytes.

## 2. METHODS

### 2.1 Production of PCSK9, CHRD and CAP1 in HEK293 FS cells

FreeStyle293FS cells (500 mL at 1.0 ×10^6^ cell/mL; Thermo Fisher) were transfected with PCSK9, CHRD and human CAP1 expression vectors (250 μg) using polyethylenimine (750 μg; Polyscience). Transfected cells were cultured at 37°C under 6% CO_2_ on an orbital shaker platform rotating at 135 rpm for 7 days before harvesting. Proteins were purified using a NiNTA purification system (HisTrap™ excel 1 mL) and size-exclusion chromatography using HiPrep™ 26/60 Sephacryl™ S-100 HR (GE Healthcare) in 10 mM Tris-HCl pH 7.5, 50 mM NaCl and 1mM CaCl_2_. Purified proteins were concentrated to 3 to 10 mg/mL (**Supplementary Figure 1**).

### 2.2 Production of P1.40 in E. coli

The expression plasmid of P1.40 carries a T7 promoter/terminator, a hexa-histidine (6xHis) tag for nickel affinity purification placed between the fusion partner (DsbC) and the Tobacco Etch Virus (TEV) protease cleavage site (ENLYFQ/G) and a gene coding for the nanobody P1.40. The optimized synthetic gene for the recombinant expression of P1.40 in *E. coli* was ordered from Eurofins. An accessory plasmid allowing the cytoplasmic expression of sulfhydryl oxidase and disulfide isomerase was used to promote the formation of disulfide bonds and the folding of P1.40 [32]. Mutagenesis of CHRD and P1.40 was performed using the QuikChange Lightning Site-Directed Mutagenesis Kit (Agilent Technologies). The integrity of the sequences of all constructs was confirmed by Sanger DNA sequencing.

Competent BL21 (DE3) pLysS were transformed with the expression and accessory plasmids and grown overnight at 37°C in 10 mL of LB medium supplemented with 50 μg/mL ampicillin and 30 μg/mL chloramphenicol. This culture was used to inoculate a Fernbach flask (1 L) of ZYP-5052 auto-medium (initial O.D._600nm_ = 0.05). ZYP-5052 medium is formulated to induce protein expression after glucose depletion. To quickly reach this step, cells were grown at 37° C (4 hours), and then at 20°C for 18h to favor soluble protein expression. After centrifugation, cells were resuspended in lysis buffer (100 mM Tris HCl pH 8, 150 mM NaCl, 5% glycerol). Benzonase (2U/mL) and MgCl_2_ (0.1 M) were then added. After 30 min at 4°C with gentle stirring, cells were lysed by two passages at 1.5 kbar on EMULSIFLEX system, centrifuged at 4°C for 45 min at 18.000 rpm, and the supernatant was recovered and filtered (0.22 μm). Proteins were loaded on a 5 ml HisTrap™HP column (1 mL/min; GE Healthcare) in a binding buffer containing 100 mM tris-HCl pH 8, 150 mM NaCl, 5% glycerol and 5 mM imidazole, and then eluted at a flow rate of 1 ml/min with a linear gradient (0 to 100%) in the same buffer containing 500 mM imidazole. Fractions containing the 6xHis-tagged fusion protein were pooled and dialyzed for 3h against a 50 mM Tris HCl pH 8, 150 mM NaCl buffer using a Spectra/Por® Dialysis Membrane (MWCO: 3500). The fusion protein was cleaved with 10% (w/w) TEV protease overnight at 4°C and the mixture was loaded on a 5 ml HisTrap™FF column (GE Healthcare). 6xHis-tag-labeled disulfide bond isomerase and TEV protease were retained, while P1.40 was recovered in flow-through. After 3h dialysis (as above), P1.40 was purified on size-exclusion chromatography (Sephacryl ® S-100 HR GE Healthcare) in 50 mM Tris HCl pH 8 and 150 mM NaCl and concentrated to 6 mg/ml (**Supplementary Figure 1**).

### 2.3 PCSK9/P1.40 and CHRD/P1.40 complexes: crystallization and structure determination of the latter

PCSK9/P1.40 and CHRD/P1.40 complexes (1:2 molar ratio molar) in 10 mM tris HCl pH 7.5, 50 mM NaCl, and 1 mM CaCl_2_ were formed overnight at 4°C, purified by size-exclusion chromatography, and analysed by electrophoresis on a blue native polyacrylamide gel system (NativePage™) (**Supplementary Figure 2A**) and purified by size-exclusion chromatography (**Supplementary Figure 2B**). Crystals were grown from 1 μl CHRD/P1.40 complex solution (5.3 mg/ml) and 1 μl precipitant from the reservoir (500 μl), by sitting drop vapor diffusion using CrysChem plates, kept at 20°C using the strategy of reverse screening [33]. A booster solution, consisting of 5 M NaCl and or 5 M NaCl with 0.2M acetic acid, was added to the reservoir to induce nucleation and the drop immediately streak seeded [34,35]. The reservoir condition used for growing the crystal consisted of 10% PEG 4000, 0.2 M imidazole malate, pH 7.0. For data collection, the crystals were cryoprotected by quick immersion into various cryoprotectant solutions [36] before flash freezing in liquid nitrogen. The data sets of several complexes were collected at synchrotron facility SOLEIL (beamlines PROXIMA-1 and PROXIMA-2, Gif-sur-Yvette, France) using both standard rotation and helical scan method [37]. About 130 samples were tested and the best crystal diffracted to 2.19 Å. X-ray diffraction data were collected from single crystals at 100 K on the Pilatus detector on beamline PROXIMA-1. The crystal belongs to the space group *P*4_1_2_1_2 with unit cell parameters of *a* = *b* = 52.3 Å and *c* = 265.2 Å. Structure deposited into the PDB Data Bank: **7ANQ**.

### 2.4 Plasmids and Mutagenesis

Human PCSK9 (full length) and CHRD (aa 449-693) cDNAs were subcloned into pIRES2-EGFP plasmid between *BglII* and *EcoRI*. Human CAP1 cDNA was cloned into pCMV plasmid between *EcoRI* and *BamHI*. All these plasmids add a hexa-histidine (6xHis) tag. Mutations were introduced in the pIRES2-EGFP-hPCSK9-WT-V5 (PCSK9-WT) plasmid by a two-step PCR approach [2]. Briefly, two-PCR fragments were generated by specific primer sets. They were digested with specific restriction enzymes and ligated. The empty vector pIRES2-EGFP-V5 was used as control. All constructs were verified by Sanger DNA sequencing. For more details on the oligonucleotides used for mutagenesis, see **Supplementary Table 1**.

The following constructs were described previously: hLDLR (pIRES2-EGFP-hLDLR-WT-V5) [38], PCSK9-Δ33-58 [39] and PCSK9-E670G [40]. Empty vector, EV (pIRES2-EGFP-V5) was used as control. PCSK9-3RA (*PCSK9-R491A-R657A-R659A*) was generated by digestion (*AgeI/PshAI*) and subsequent ligation of resulting fragments from PCSK9-R491A and PCSK9-R657A/R659A constructs. PCSK9-Δ33-58/3RA was generated by digestion (*AgeI/SacI*) and subsequent ligation of resulting fragments from PCSK9-Δ33-58 and PCSK9-3RA constructs. The following constructs were described previously: hLDLR (pIRES2-EGFP-hLDLR-WT-V5) [38], PCSK9-Δ33-58 [39] and PCSK9-E670G [40]. All constructs were verified by Sanger DNA sequencing.

### 2.5 Cell culture and analysis

HepG2-*PCSK9*^*-/-*^(CRISPR/cas9-PCSK9 [40]), IHH and Huh7 cells were grown in EMEM medium (Wisent) supplemented with 10 % fetal bovine serum, FBS (GIBCO BRL) and transfected using Fugene HD reagent (Promega). HEK293 cells were cultured in DMEM medium (Wisent) plus 10 % FBS and transfected using jetPRIME (Polyplus) reagent. All cells were kept at 37°C under 5% CO_2_ and transfected at 50-70 % confluency according to the manufacturer’s instructions. 24h post-transfection, the cells were pre-incubated in serum-free medium, SFM for 1 h, then incubated 18h with conditioned media. Finally, the media and cells were collected for analysis. The media were centrifuged (5 min at 12,000 rpm, 4°C), supplemented with 50 mM sodium fluoride (NaF) (BioShop) and protease inhibitor cocktail, PIC (Roche Applied Science) and stored at -80°C until analysis. The cells were washed with ice-cold PBS, collected, and lysed in RIPA buffer (50 mM Tris-HCl pH 7.5, 150 mM NaCl, 2 mM EDTA, 50 mM NaF, 1% Nonidet P-40, 0.5 % sodium deoxycholate, 0.1 % SDS, and PIC). The resulting total cell lysates and media were analysed by Western blotting, imaged using ChemiDoc MP imaging system (Bio-Rad) and the protein bands quantified by Image LabTM Software (Bio-Rad) as previously described [40]. The following antibodies were used in this study: anti-V5-epitope (1:5,000; Sigma), monoclonal antibody to detect V5-tagged proteins (PCSK9 and LDLR), anti-Flag (1:2,000; Sigma); anti-HA (1:4,000; Sigma), anti-hLDLR (1:1,000; R&D system), anti-hPCSK9 (1:2,000; [14]), anti-β-actin (1:3000; Sigma), anti-CAP1 (1:2,000; Santa Cruz), anti-Caveolin-1 (1:1,000; abcam), anti-Clathrin Heavy Chain (1:1,000; Invitrogen); anti-goat-conjugated Alexa-488 (1:500; Molecular probes, Invitrogen), and anti-rabbit (1:10,000; Amersham), anti-goat (1:10,000; Sigma) and anti-mouse (1:10,000; VWR)-conjugated to horseradish peroxidase HRP.

### 2.6 HEK293 cells conditioned media production and media swap

PCSK9 conditioned media were produced following HEK293 cells transfection with PCSK9 constructs as described above. The media were collected 48h post-transfection, centrifuged 5 min (12,000 rpm/4°C), quantified by ELISA, and stored at -80°C until use [41]. For media swap, cells were pre-incubated in SFM for 1 h, followed by conditioned media swap for 18h after which both cells and media were collected for analysis. P1.40 and mutant P1.40-R105A nanobodies were produced at 6 mg/ml and 1.6 mg/ml concentrations respectively and used at 1 μg/ml final concentration. Purified nanobodies were preincubated 1h with conditioned media then added to the cells for 18h incubation.

### 2.7 Immunocytochemistry and Dil-LDL uptake assay

For immunocytochemistry experiments, HepG2-*PCSK9*^−/-^cells (0.5×10^5^ cells/well) were plated on poly-L-lysine-coated round microscope coverslips that were placed in a 24-well cell culture plate. Cells were then treated as required for each experiment (siRNA transfection, protein overexpression, PCSK9 incubation). To analyze plasma membrane LDLR and CAP1 expression, the cells were washed twice with PBS and fixed with a solution of 4 % paraformaldehyde in PBS (10 min). After blocking with PBS BSA 2% (1h), samples were incubated at 4°C overnight with the respective primary antibodies (LDLR, R&D Systems, 1:200 and CAP1, Santa Cruz, 1:1,000). Cells were then washed with PBS and incubated with the appropriate fluorescent secondary antibody (all from Thermo Fisher, 1/500) for 1h at room temperature. To analyze total LDLR and CAP1 expression, cells were treated with the same protocol but were permeabilized with 0.2% TritonX-100 in PBS for 5 min prior blocking. To test LDLR functionality, cells were incubated for 2h at 37°C with 5 μg/ml DiI-LDL (1,1’-dioctadecyl-3,3,3’,3’-tetramethyl-indocarbocyanine perchlorate, Cedarlane/Kalen Biomedical) in SFM media before fixation. Coverslips were mounted on a glass slide with ProLong Gold antifade reagent with DAPI (Thermo Fisher). Samples were visualized using Plan-Neofluar 40x/0.60 or Plan-Apochromat 63x 1.4 oil immersion objectives of LSM-710 confocal laser-scanning microscope (Carl Zeiss) with sequential excitation and capture image acquisition with a digital camera. Images were processed with ZEN software. Image analysis to quantify the fluorescence intensities was accomplished using Volocity®6.0. Dil-LDL uptake assay was achieved as previously described [40].

### 2.8 PCSK9 and MHC-I

#### Database search for the motif RXE

A python code was written to analyze all non-redundant protein sequences from NCBI’s BLAST database (with entries from GenPept, Swissprot, PIR, PDF, PDB, and RefSeq; downloaded June 14, 2021 at: https://ftp.ncbi.nlm.nih.gov/blast/db/). Briefly, all human non-redundant sequences with one or more occurrences of the motif RXE (X = any residue) were filtered to a tab separated values file (.tsv) corresponding to the following fields: the fasta header, the protein sequence, the list of motifs that were found and, the total count of motifs for the sequence. A multi-fasta file was generated using all listed motifs. Next, using WebLogo3.7 [42].

#### Molecular modeling of PCSK9 / HLA-I complex

PCSK9 (Pdb ID: 2P4E) and HLA-I (Pdb ID: 6TRN) structures were docked using GRAMM-X web server (http://vakser.compbio.ku.edu/resources/gramm/grammx/, [43] respectively, as receptor and ligand. All available chains in PDB were used and the residues R44 and E46 of HLA-I, comprised in the motif RXE, were assumed as potential interface residues. The interface residues of the docking model were subjected to energy minimization using GROMOS96 43B1 [44] implementation of Swiss-PdbViewer (100 steepest descent steps, 1 conjugate gradient step, 100 steepest descent steps). The quality of the final model was evaluated using PROCHECK [45].

## 3. RESULTS

### 3.1 Structural analysis of the complex CHRD-P1.40 nanobody

Binding of P1.40 to the CHRD inhibits the PCSK9-mediated degradation of the LDLR [27,28]. We first assessed the affinity of full-length PCSK9 or the CHRD alone for biotinylated P1.40 by Bio-Layer Interferometry (BLI; **Figure 1A**). A 1:1 binding model describing a single exponential function for both complexes was used to calculate kinetic constants demonstrating that the association kinetics of P1.40 to the CHRD is 238 times faster (k_a_ = 3.8 ×10^5^ ± 0.1×10^5^ M^−1^s^−1^) compared to full-length PCSK9 (k_a_ = 1.6×10^3^ ± 0.1×10^3^ M^−1^s^−1^), whereas the dissociation kinetics are similar for the two complexes. Thus, the estimated K_D_ values of 5.6 nM and 90.3 pM, respectively, indicate that P1.40 has a 62-fold greater affinity for the CHRD alone compared to full-length PCSK9.

**Figure 1:**
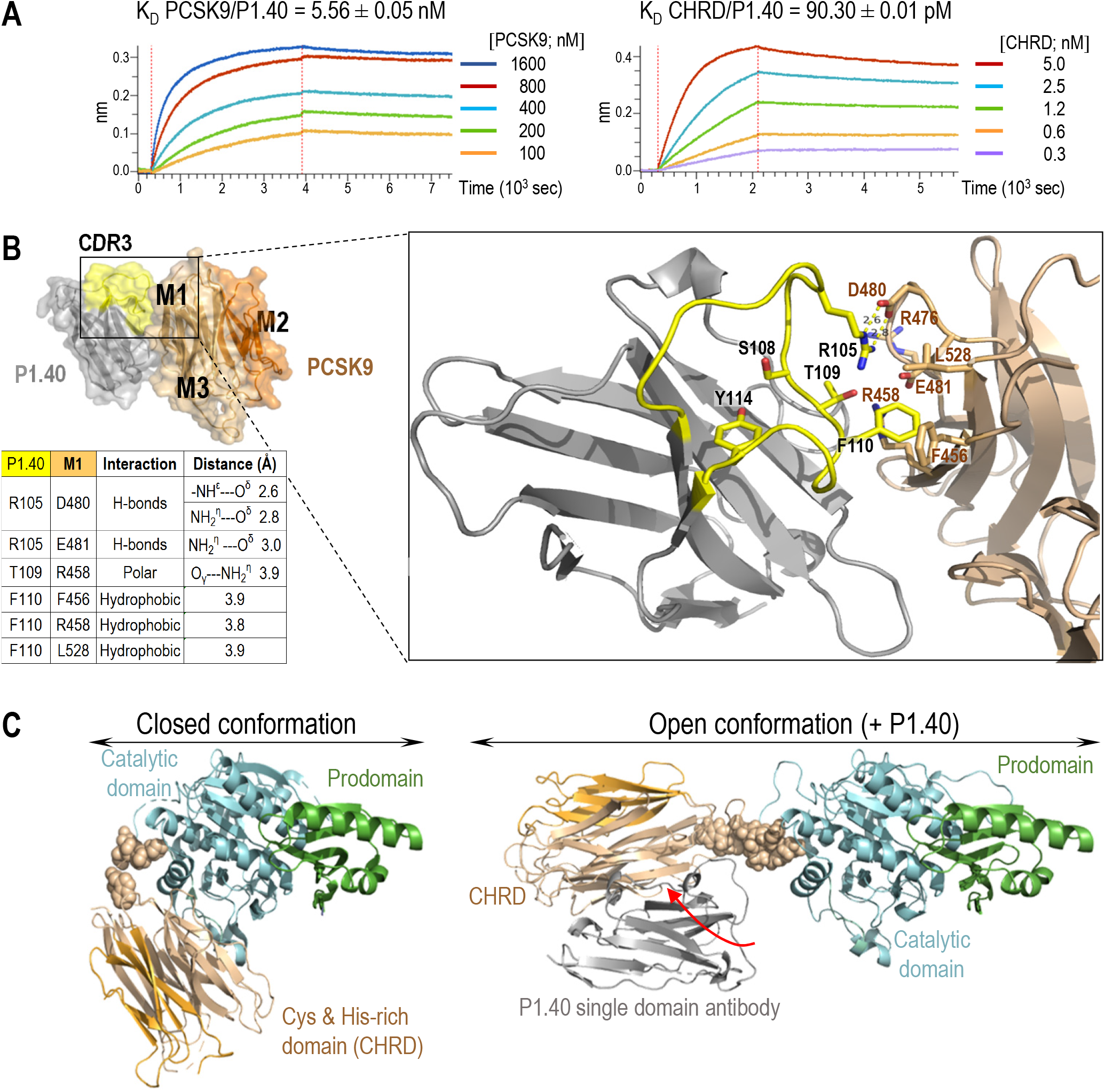
Characterization of PCSK9 and P1.40 interaction. **(A)** Sensorgrams show real-time BLI assays in which kinetic constants were measured at pH 7.5 for PCSK9/P1.40 and CHRD/P1.40 complexes using 0.25 μg/μL of immobilized biotinylated P1.40. PCSK9/P1.40 complex: association time 3600 s, k_a_ = 1.6×10^3^ ± 0.1×10^3^ M^−1^s^−1^; dissociation time 3600 s; k_diss_ = 9.0 ×10^−6^ ± 0.1×10^−6^ s^−1^. CHRD/P1.40 complex: association time 1800 s, k_a_ = 3.8 ×10^5^ ± 0.1×10^5^ M^−1^s^−1^, dissociation time 3600 s; k_diss_ = 3.5×10^−5^ ± 0.1×10^−5^ s^−1^. (**B**) The surface view of the CHRD/P1.40 crystal (PDB 7ANQ) obtained at pH 7.5 shows the M1 and M3 (brown) and the M2 (light orange) subdomains of the CHRD, and P1.40 (grey). The highly variable loop “complementary-determining region 3” of P1.40 (yellow; aa 105-122; CDR3) contributes to antigen recognition. A zoom view shows the main interactions between P1.40 (yellow sticks) and M1 (brown sticks) residues. P1.40 Arg_105_ establishes three salt bridges (COO^−^---NH ^+^ [74]) with M1: two with Asp_480_ and one with Glu_481_. In addition, hydrogen bonding occurs in P1.40, between Arg_105_ and the OH group of Thr_109_ (3.3 Å, *not shown*), thereby maintaining a favorable orientation when Arg_105_ interacts with Asp_480_ in M1. Thr_109_ also interacts with Arg_458_ in M1. In addition, P1.40 Phe_110_ is sandwiched *via* Van der Waals interactions between M1 Phe_456_, Arg_458_ and Leu_528_. Finally, the critical role of Tyr_114_, which makes no contact with CHRD residues, is probably to lock the conformation of the long P1.40 loop by an intramolecular hydrogen bond between its OH and the carbonyl of the peptide bond of Ser_108_. (**C**) Best DADIMODO atomic models for PSCK9 alone in a “closed” conformation and in complex with P1.40 in an “open” conformation. In both conformations, the pro- and catalytic domains are shown in the same orientation. The flexible linker (spheres; aa 422-452) between the catalytic domain and the CHRD allows the rotation of the latter upon P1.40 interaction (red arrow).

The crystal of the P1.40-CHRD complex belongs to the tetragonal space group P4_1_2_1_2 with the cell parameters a = b = 52.3 Å, c = 265.2 Å and α = β = γ = 90°. The diffraction limit of 2.2 Å was recorded for the best data set collected (**Supplementary Table 2**). The major surface contact between P1.40 and the CHRD is an area of 850 Å^2^ (Protein Interfaces, Surfaces and Assemblies program [46]) that essentially involves Arg_105_, Thr_109_ and Phe_110_ (yellow sticks) of P1.40 and Phe_456_, Arg_458_, Arg_476_, Asp_480_, Glu_481_ and Leu_528_ (brown sticks) of the M1 subdomain of the CHRD (**Figure 1B**). Additional interactions between P1.40 (Arg_43_, Glu_44_, Glu_46_, Asp_62_, Asp_89_ and Arg_111_) and M1 and M3 subdomains (Thr_459_, Trp_461_, His_464_ Arg_476_ and Asp_651_) were also observed (**Supplementary Figure 3**).

Some of the identified M1 residues (Phe_456_, Arg_476_, Asp_480_) were substituted to Ala and the mutants were produced in HEK293 cells to investigate their contribution to P1.40 recognition. Cell binding assays using P1.40 expressed on yeast cells revealed a dramatic affinity decrease only for the D480A mutant (142-fold) and a 12- and 17-fold reduction for the F456A and R476A mutants, respectively (**Supplementary Figure 4A**). Thus, the P1.40 Arg_105_ and CHRD Asp_480_ interaction plays a critical role in the complex stability. In contrast, pairwise interactions between Asp_62_ and Phe_110_ of P1.40 and Arg_476_ and Phe_456_ of the CHRD, respectively, contribute less to complex stability (**Figure 1B** and **Supplementary Figure 3B**).

We corroborated the above data using a Deep Mutational Scanning approach for residues 97 to 116 that cover the junction between a conserved structural segment and a highly variable loop (residues 105 to 122) known to contribute to antigen recognition. Yeast cells with a preserved expression of cell surface HA-tagged P1.40 mutants were selected by FACS and sorted according to their ability to bind the 6xHis-tagged CHRD (**Supplementary Figure 4B**). As expected, the strongest losses of binding were observed for amino acid changes in the highly variable loop at Arg_105_, Thr_109_, Phe_110_ and Tyr_114_. For these residues, Ala-substitutions led to marked losses of affinity (**Supplementary Figure 4C**). Finally, cell validation was achieved in human hepatic HepG2 cells, in which *PCSK9* expression was silenced by CRISPR-Cas9 (HepG2-*PCSK9*^−/-^) [40]. Compared to P1.40, its R105A mutant no longer inhibits the ability of extracellular PCSK9 to enhance the degradation of the LDLR [27] (**Supplementary Figure 5A**). In agreement, the PCSK9 natural variant D480N, while having an activity on LDLR comparable to that of wild-type (WT) PCSK9, is no longer sensitive to P1.40 (**Supplementary Figure 5B**).

To explain the higher affinity of P1.40 for the CHRD alone *versus* full-length PCSK9, the 3D structure of the CHRD/P1.40 (PDB code 7ANQ) was superimposed on that of PCSK9 (PDB code 2P4E) with a PyMOL software (https://pymol.org/2/). This overlay highlights that a large part of the CHRD surface involved in the P1.40 interaction is hidden in the crystal structure of PCSK9. Complex formation would thus require opening of the PCSK9 “closed conformation” to expose to solvent the surface involving P1.40 interactions. Such movement should be possible as the two domains of PCSK9, the catalytic domain and CHRD, are connected through a 19-residue linker (Lys_421_-Asn_439_) essential for LDLR degradation [9]. To validate this hypothesis, size exclusion chromatography followed by Small Angle X-ray Scattering (SAXS) were performed with PCSK9 alone or in complex with P1.40 to detect large conformational shifts in solution (**Supplementary Table 3**; **Supplementary Figure 6**). We demonstrated by size exclusion chromatography that in solution PCSK9 is monomeric and that the PCSK9/P1.40 complex remains stable and does not dissociate (**Supplementary Figure 6A**). The information deduced from the average experimental SAXS curves (**Supplementary Figure 6B-D**) allowed us to estimate a molecular weight of PCSK9 alone (70.6 kDa) and of the PCSK9/P1.40 complex (88.3 kDa), indicating that a single protein P1.40 (14.4 kDa) binds to PCSK9 (72.5 kDa) in solution (**Supplementary Figure 6E**) and that the PCSK9/P1.40 complex is more extended than PCSK9 alone (**Supplementary Figure 6F**). To understand the conformational changes induced by P1.40 binding to PCSK9, atomic models in “closed” and “open” conformations were generated using the MODELLER software [47] from the 3D structure of PCSK9 (PDB code 2P4E) and the 3D structure of PCSK9 and CHRD-P1.40 (PDB code 2P4E + 7ANQ). These latter models in “open” and “closed” conformations were then used to generate atomic models under the constraints of average experimental SAXS curves using the DADIMODO software [48]. Our data show that the formation of the PCSK9/P1.40 complex implies a significant movement of the CHRD domain relative to the catalytic domain that leads to a transition from a “closed” to an “open” conformation of PCSK9 (the best DADIMODO models in **Figure 1C**). This rationalizes the inhibitory effect of P1.40 on the PCSK9 activity on the LDLR, as P1.40 would primarily bind the CHRD in the “open” PCSK9 conformation and interact more effectively with the CHRD alone. Indeed, the presence of two PCSK9 conformations that bind the LDLR was previously predicted from the first reported crystal structure of PCSK9 [9].

### 3.2 Secreted CAP1 enhances PCSK9 activity on the LDLR *via* endocytosis in clathrin-coated vesicles

The mechanism by which CAP1 enhances PCSK9-mediated LDLR degradation is not known. We investigated CAP1 function in HepG2-*PCSK9*^−/-^ cells [40] that were incubated with conditioned media from HEK293 cells (media swap) which expressed an empty vector or PCSK9-V5 [23,40], or were supplemented with purified recombinant PCSK9 (rPCSK9) [38]. While the loss of endogenous CAP1 (siCAP1) in HepG2-*PCSK9*^−/-^ cells reduced extracellular PCSK9 activity it did not abrogate its function (**Figure 2A**). Unexpectedly, cytosolic CAP1 was abundantly secreted into the media of HepG2-*PCSK9*^−/-^ cells (siCTL; **Figure 2A**). Thus, CAP1 is not only a cell-surface associated protein but could potentially bind PCSK9 extracellularly. The functional activity of PCSK9 on LDLR in absence of endogenous CAP1 was further confirmed by immunocytochemical analysis of LDLR and DiI-LDL uptake of non-permeabilized cells (**Figure 2B**). PCSK9 reduced the cell surface levels of both LDLR and CAP1 (**Figure 2B**), suggesting that the complex LDLR-PCSK9-CAP1 is targeted for degradation. Interestingly, to get similar reduction in LDLR levels as those achieved by PCSK9 secreted from HEK293 cells (-65% to -74%) we had to incubate HepG2-*PCSK9*^−/-^ cells with ∽10-fold higher levels of rPCSK9 (**Figure 2A**), as previously reported [38]. Thus, we hypothesized that the stronger LDLR reduction by PCSK9-V5 as compared to rPCSK9 may be due to the endogenous expression of CAP1 in HEK293 cells, in which PCSK9-V5 was produced. Accordingly, we performed the same experiment by producing PCSK9-V5 in the medium of HEK293 cells pre-treated with CAP1 siRNA and tested its effect on HepG2-*PCSK9*^−/-^ cells lacking endogenous CAP1 (**Figure 2C**). Under these conditions, PCSK9-V5 activity was significantly reduced, revealing that CAP1 is not essential but contributes to enhancing PCSK9 activity on the LDLR.

**Figure 2:**
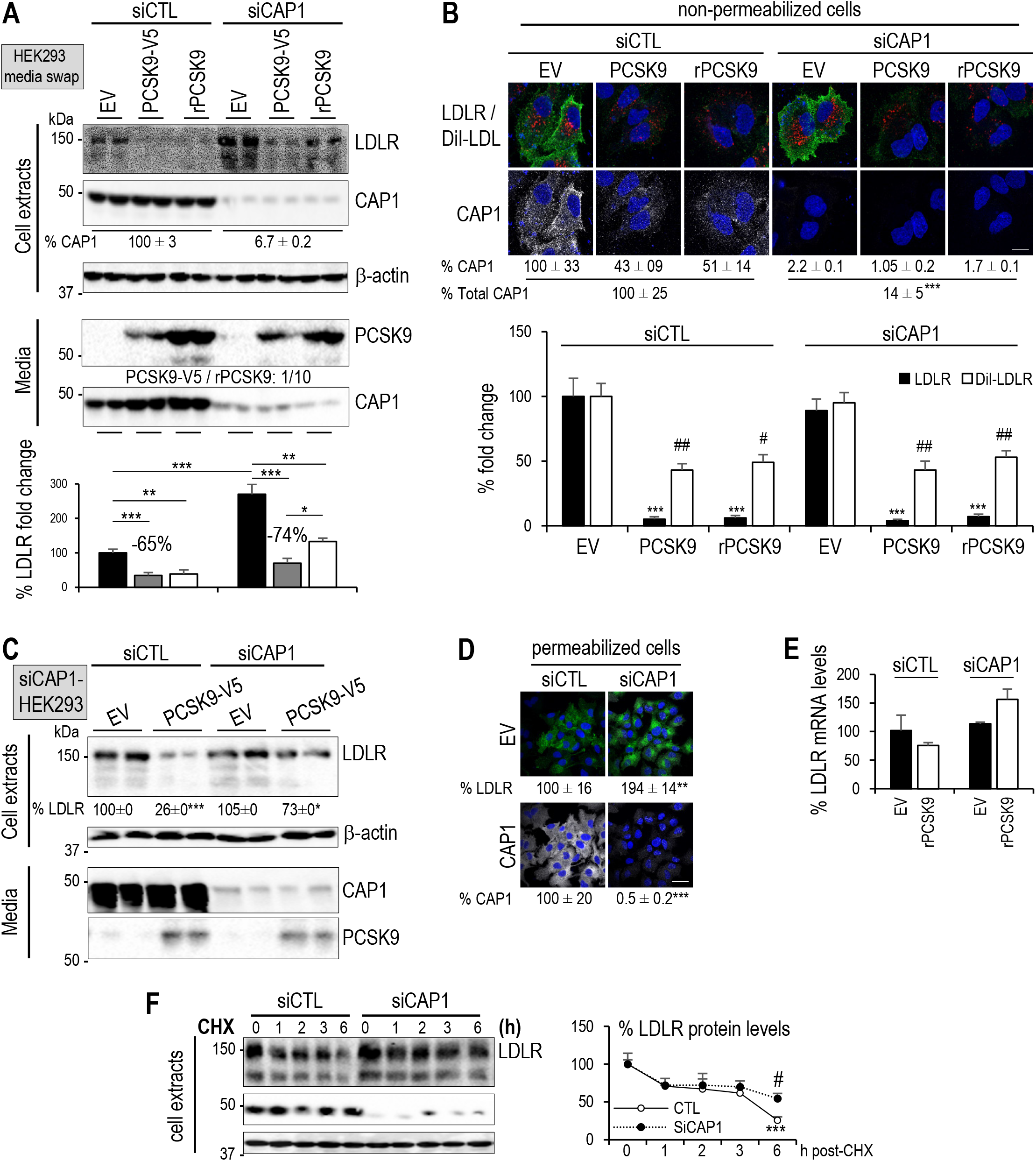
Effect of CAP1, caveolin-1 and clathrin heavy chain in PCSK9-mediated LDLR degradation. (**A**) HepG2-*PCSK9*^−/-^ cells were transfected with control (siCTL) or CAP1 (siCAP1) siRNAs and the next day incubated with conditioned media from HEK293 cells expressing an empty vector (EV) or PCSK9-V5 (0.3 μg/ml). A third subset of HepG2-*PCSK9*^−/-^ cells was incubated with recombinant PCSK9 (rPCSK9; 3 μg/ml) added to a fresh medium. After 18 h, cell extracts and media were analyzed by Western blot using appropriate antibodies. LDLR levels were normalized to those in siCTL/EV and set to 100. (**B**) HepG2-*PCSK9*^−/-^ cells were transfected with siCTL or siCAP1 and incubated with conditioned media from HEK293 cells expressing empty vector (EV) or PCSK9-V5 (0.3 μg/ml). A third subset was incubated recombinant PCSK9 (rPCSK9; 3 μg/ml) added to fresh medium. Non-permeabilized cells were incubated with DiI-LDL for 2h, fixed and analyzed by immunocytochemistry for LDLR (green; black bars) and DiI-LDL internalization (red; white bars). In contrast, CAP1 levels (white; indicated values) were assessed in permeabilized cells. All cells were stained with DAPI (blue). Representative images are shown. Mean ± SEM are given for four independent experiments. Scale bar is 20 μm. (**C**) LDLR and secreted CAP1 protein levels were analyzed in HepG2-*PCSK9*^*-/-*^ cells that underwent the same treatment as in (A), except that HEK293 cells were treated with CAP1 siRNA before transfection and medium swap. (**D**) LDLR immunocytochemistry (green) was performed on fixed and permeabilized cells treated as in (A) and stained with DAPI (blue). Scale bar is 50 μm. **(E**) LDLR mRNA levels were assessed in the HepG2-*PCSK9*^−/-^ cells that were analyzed in (A). **(F**) HepG2-*PCSK9*^*-/-*^ cells were transfected with siCTL or siCAP1 for 24h and then incubated with 40 mg/mL cycloheximide (CHX) to follow the LDLR degradation within 6 h. LDLR protein levels were normalized to those in HepG2-*PCSK9*^*-/-*^ cells treated with siCTL and set to 100. Representative data and images and mean ± SEM are shown for ≥3 independent experiments. *P* values for LDLR and CAP1 (*, *P*<0.05; **, *P*<0.01; ***, *P*<0.001) or DiI-LDL (^#^, *P*<0.05; ^##^, *P*<0.01) were obtained by two-way ANOVA coupled to Tukey’s multiple comparisons tests (B) and by Welch’s t-test (D).

Unexpectedly, while not affecting cell surface LDLR levels (**Figure 2B**), siCAP1 treatment of HepG2-*PCSK9*^−/-^ cells (EV) enhanced total LDLR levels by ∽2.5-fold in the absence of PCSK9 (**Figure 2A**), as supported by immunocytochemical analysis of permeabilized cells (**Figure 2D**), suggesting a PCSK9-independent intracellular mechanism. Importantly, siCAP1 treatment did not affect LDLR mRNA levels (**Figure 2E**). To begin probing the cause of siCAP1-induced enhanced total LDLR levels, we incubated HepG2-*PCSK9*^−/-^ cells with cycloheximide to block mRNA translation [49]. The data revealed that the effect of CAP1 siRNA on endogenous LDLR protein occurs at least in part *via* its stabilization, with a significant ∽2-fold increase after 6h of incubation (**Figure 2F**). This may be related to the ability CAP1 to accelerate actin turnover dynamics [50].

PCSK9 was reported to enhance the degradation of the LDLR following endocytosis into clathrin heavy chain (CHC)-coated vesicles [51]. However, it was also suggested that the PCSK9/LDLR complex is directed to caveolin-coated endosomes in the presence of CAP1 [30]. We thus verified the effect of CAP1 on PCSK9-mediated LDLR degradation in the presence (siCTL) and absence of clathrin-heavy chain (siCHC) or caveolin-1 (siCav1), the main component of caveoli in hepatocytes [52]. Western blotting showed that, in the presence of endogenous CAP1, siCHC, but not siCav1, completely blocked the activity of extracellular PCSK9 (**Figure 3A**). Immunocytochemistry of non-permeabilized cells confirmed the lack of effect of siCAP1 on cell surface LDLR levels, but also clearly showed that siCHC prevented PCSK9 activity on the LDLR, but not siCav1 (**Figure 3B**). Thus, in our hands even in the presence of endogenous CAP1, PCSK9-LDLR complexes are degraded following their internalization into clathrin-coated vesicles, but not caveoli.

**Figure 3:**
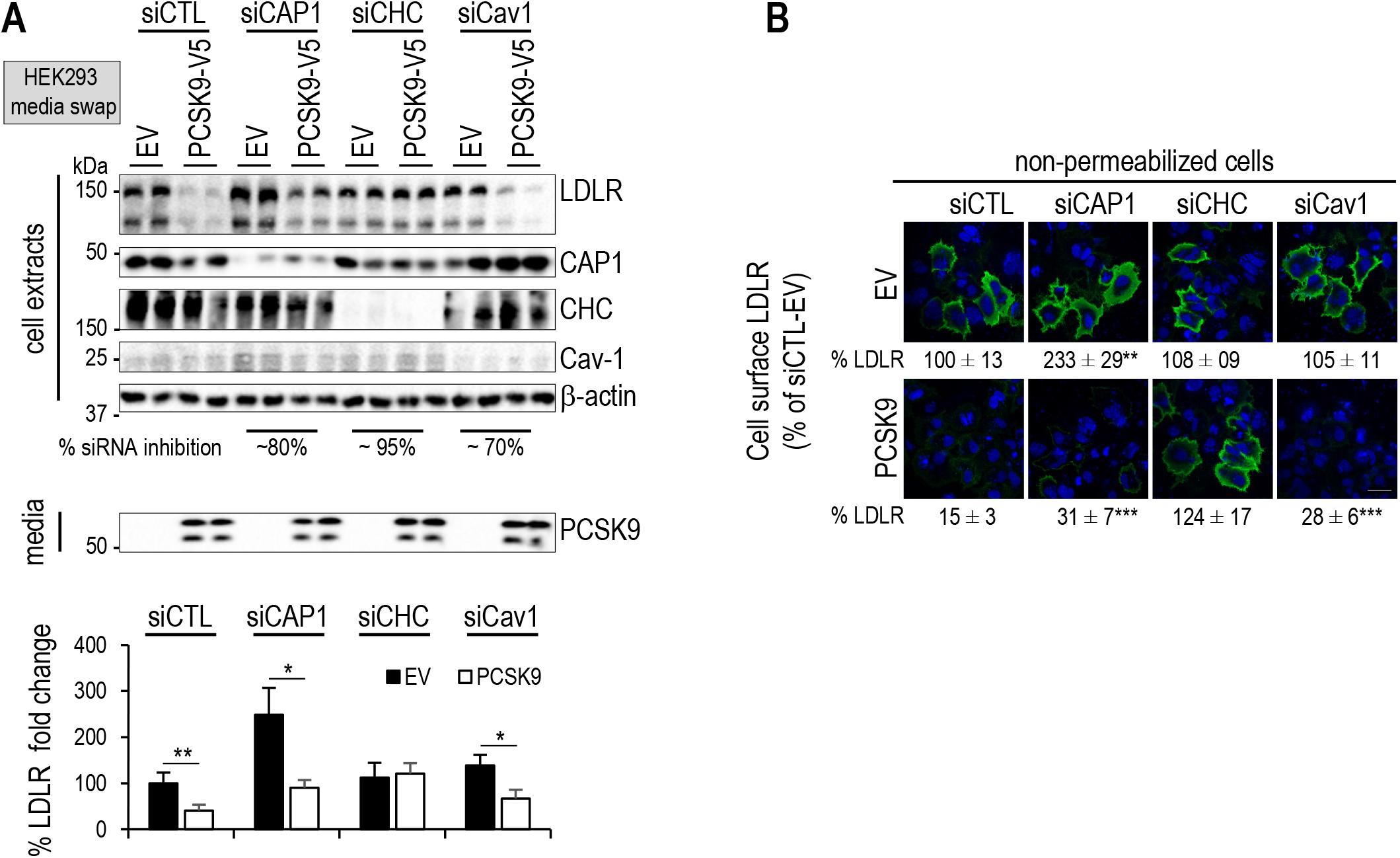
Effect of CAP1 silencing on LDLR stability and PCSK9-mediated LDLR degradation. **(A)**HepG2-*PCSK9*^−/-^ cells were transfected with CTL siRNA, siRNAs against CAP1, clathrin heavy chain (siCHC) or caveolin-1 (siCav1). The average % silencing achieved by each siRNA is shown. The cells were then incubated with conditioned media from HEK293 cells expressing an empty vector (EV) or PCSK9-V5 (0.3 μg/ml). LDLR intensity was normalized as in (Figure 2A). (**B**) LDLR immunocytochemistry (green) was performed in fixed non-permeabilized cells treated as in (Figure 2B) and stained with DAPI (blue). Scale bar is 20 μm. Representative data and images and mean ± SEM are shown for ≥ 3 independent experiments. All *P* values (*, *P*<0.05; **, *P*<0.01; ***, *P*<0.001) were obtained by two-way ANOVA coupled to Tukey’s multiple comparisons tests.

### 3.3 CAP1 and P1.40 bind the M1 and M3 domains of the CHRD at different sites

The available 3D-structures of the CHRD [9,10] prompted us to test the effect of Ala-substitutions of three exposed residues in M1 (Arg_491_,) and M3 (Arg_657_ and Arg_659_) subdomains on PCSK9 activity (triple CHRD mutant; 3RA). Although these substitutions had no effect on the CHRD-P1.40 interaction (**Supplementary Figure 4A**), they dramatically reduced the activity of PCSK9 on the LDLR (**Figure 4A**), as well as CHRD binding to biotinylated CAP1, as estimated by BLI (**Figure 4B**), suggesting that the targeted residues contribute at least in part to CAP1 binding. Altogether, these data indicate that CAP1 and P1.40 bind different sites in the M1 and M3 subdomains of the CHRD. Analysis by BLI of CAP1 binding to biotinylated PCSK9 revealed a K_D_ value of 1.36 μM that agrees with the previously reported one [30]. A similar K_D_ of 1.26 μM is found for CAP1 binding to the biotinylated CHRD alone (**Figure 4C**). Indeed, similar kinetic constants were obtained for PCSK9 and the CHRD alone: k_ass_ = 6.9 ×10^2^ ± 0.1×10^2^ M^−1^s^−1^ and k_ass_ = 5.6 ×10^2^ ± 0.1×10^2^ M^−1^s^−1^; k_diss_ = 9.4 ×10^−4^ ± 0.1×10^−4^ s^−1^ and k_diss_ = 7.0 ×10^−4^ ± 0.1×10^−4^ s^−1^, respectively, demonstrating that CAP1 binds with similar affinity both PCSK9 and the CHRD (**Figure 4C**).

**Figure 4:**
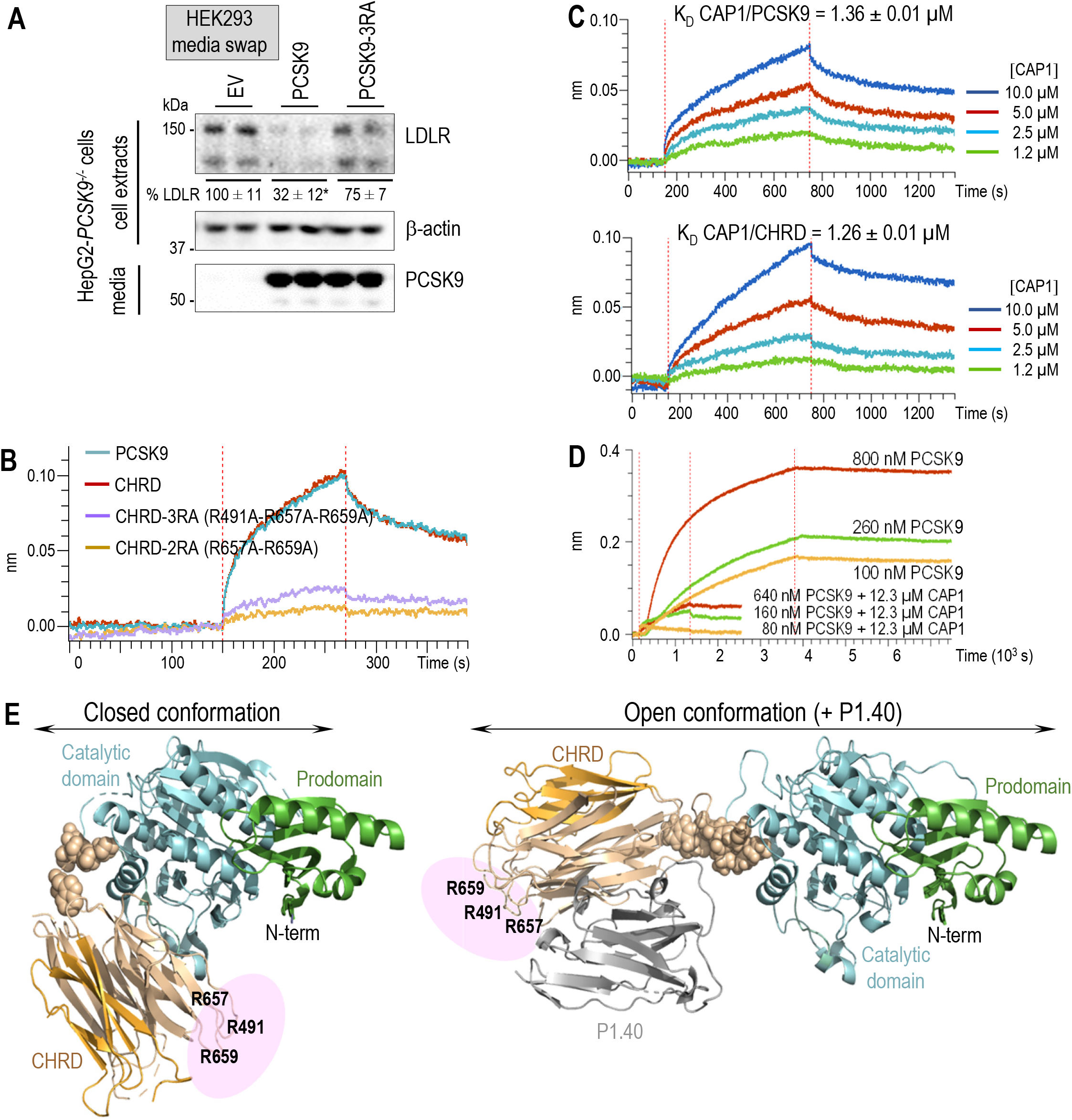
CAP1 and P1.40 bind the M1 and M3 domains of the CHRD at different sites. (**A**) HepG2-*PCSK9*^−/-^ cells were incubated with conditioned media from HEK293 cells expressing empty vector (EV), PCSK9 or PCSK9-3RA (0.3 μg/ml) for 18 h. Cell extracts and media were analyzed by Western blot. Data represent means ± SEM of two independent experiments. *P* values for LDLR (*, *P*<0.05) were obtained by two-way ANOVA coupled to Tukey’s multiple comparisons tests. (**B**) Sensorgrams show real-time BLI assays in which kinetic constants were measured at pH 7.5 for CAP1/PCSK9 and CAP1/CHRD complexes using 0.2 μg/ml of biotinylated PCSK9 or biotinylated CHRD. Association and dissociation time were 600 s for both. CAP1/PCSK9 complex: k_a_ = 6.9×10^2^ ± 0.1×10^2^ M^−1^s^−1^ and k_diss_ = 9.4 ×10^−4^ ± 0.1×10^−4^ s^−1^. CAP1/CHRD complex: k_a_ = 5.6×10^2^ ± 0.1×10^2^ M^−1^s^−1^ and k_diss_ = 7.0 ×10^−4^ ± 0.1×10^−4^ s^−1^. (**C**) Binding of PCSK9, CHRD, CHRD-3RA or CHRD-2RA (all at 10 μM) to immobilized biotinylated CAP1 (2 μg/μL) was analyzed by BLI. The association and dissociation time were 120 s. (**D**) Sensorgrams for PCSK9 alone (association time 3600 s and dissociation time 3600 s) or pre-formed PCSK9+CAP1 complexes (association time 1200 s and dissociation time 1200 s) to immobilized biotinylated P1.40 (0.25 μg/μL). (**E**) Schematic representation of the putative binding site of CAP1 (pink) to the CHRD based on the best DADIMODO atomic models for PCSK9 alone (closed conformation) or in complex with P1.40 (open conformation).

Next, we analyzed by BLI the binding of increasing concentrations of PCSK9 that was not (upper curves) or was (lower curves) pre-incubated with a 19, 77 or 154-fold excess of CAP1 to biotinylated P1.40 (**Figure 4D**). The binding curves revealed that, in the presence of an excess of CAP1, PCSK9 no longer bound P1.40. In conclusion, CAP1 seems to be able to bind both “closed” and “open” conformations of PCSK9, even in presence of P1.40. However, by stabilizing the closed conformation of PCSK9, CAP1 likely prevents the access of P1.40 to its binding site on PCSK9 (**Figure 4E**).

### 3.4 Overexpressed CAP1 prevents P1.40 inhibition of cellular PCSK9 activity on the LDLR

*In vitro*, P1.40 binds PCSK9 (**Figure 1A**; K_D_ 5.56 nM) with a ∽250-fold higher affinity than CAP1 (**Figure 4B**; K_D_ 1.36 μM). We thus tested LDLR regulation by PCSK9 upon CAP1 overexpression in HepG2-*PCSK9*^−/-^ cells that were incubated with the media of HEK293 cells expressing PCSK9 alone or with exogenously added P1.40. As expected from the BLI analysis (**Figure 4D**) and SAXS’s atomic models (**Figure 4E**) overexpression of CAP1 abrogated the inhibitory effect of extracellular P1.40 (**Figure 5A**), an effect that was confirmed by immunocytochemistry (**Figure 5B**) and in two other hepatocyte cell lines, i.e., IHH and Huh7 cells (**Supplementary Figure 7A**,**B**). Since Asp_480_ is critical for PCSK9 binding to P1.40 Arg_105_ (**Figure 1B**), we evaluated the impact of D480N substitution on PCSK9 inhibition by P1.40 in the presence or absence of overexpressed CAP1. Although PCSK9-D480N was as potent as WT PCSK9 on the LDLR, this mutation abrogated the inhibitory properties of P1.40, irrespective of the presence or absence of CAP1 (**Supplementary Figure 7C**). Thus, the hotspot M1 Asp_480_ needed for PCSK9-P1.40 interaction has no impact on PCSK9-CAP1 binding. We conclude that CAP1 may favor the closed configuration of PCSK9 thereby preventing P1.40 binding to the CHRD and hence PCSK9 inhibition (**Figure 4E**).

**Figure 5:**
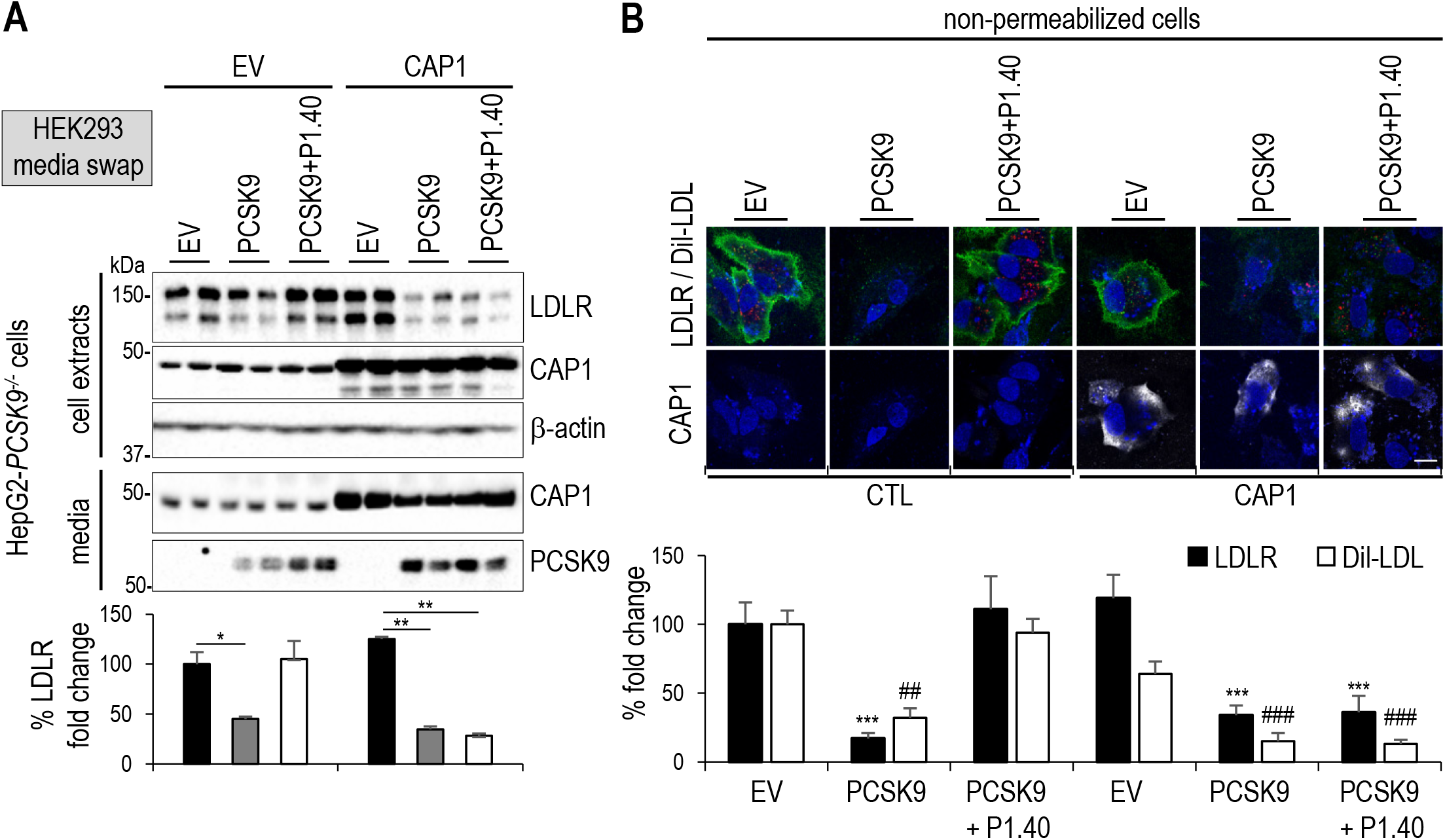
Effect of CAP1 overexpression on PCSK9 activity. (**A**) HepG2-*PCSK9*^*-/-*^ cells were transfected with a vector either empty (EV) or expressing CAP1. After 24 h, cells were incubated with conditioned media from HEK293 cells expressing an empty vector (EV) or PCSK9 (0.3 μg/ml), in the presence or absence of P1.40 (1 μg/mL) for 18 h. Cell extracts and media were analyzed by Western blot and normalized LDLR levels are shown. (**B**) HepG2-*PCSK9*^−/-^ cells were transfected with an empty (EV) or CAP1-expressing vector and treated as in (A). Cells were then incubated for 2h with DiI-LDL prior to immunocytochemistry of cell surface LDLR (green), DiI-LDL (red) and CAP1 (white) under non-permeabilized conditions and stained with DAPI (blue). Representative images and mean ± SEM from three independent experiments are shown. Scale bar is 20 μm. *P* values for LDLR (*, *P*<0.05; **, *P*<0.01; ***, *P*<0.001) or DiI-LDL (^##^, *P*<0.01; ^###^, *P*<0.001) were obtained by two-way ANOVA coupled to Tukey’s multiple comparisons tests.

Even though PCSK9-3RA exhibited a substantial loss of activity on the LDLR in the presence of endogenous levels of CAP1 (**Figure 4A**), overexpressed CAP1 restores the mutant’s activity to WT-PCSK9 levels (**Supplementary Figure 7D**). This indicated that CAP1 could also interact with predicted other M1 and M3 CHRD residues (e.g., Lys_494_, Ser_642_, Asp_660_ and Ser_662_) [30]. In addition, because the unstructured N-terminal region of the PCSK9 prodomain (aa 31-58) [53,54] is rich in acidic residues that are known to reduce PCSK9 activity [54], we hypothesized that CAP1 binding to some of these residues may neutralize them and may further explain its capacity to increase PCSK9-mediated LDLR degradation. We thus tested the ability of CAP1 to stimulate the activity of the truncated PCSK9-Δ33-58 [39]. Amazingly, upon CAP1 overexpression, P1.40 still inhibited the function of PCSK9-Δ33-58, indicating that it is in an open conformation (**Figure 6A**). Co-immunoprecipitations of HA-tagged CAP1 with V5-tagged PCSK9 or PCSK9-Δ33-58 show that P1.40 reduces CAP1 binding to WT-PCSK9, but not PCSK9-Δ33-58, which exhibits lower binding affinity to CAP1 (**Figure 6B**). We are thus proposing the existence of an interaction between CAP1 and some acidic residues of the prodomain (Asp_35_, Glu_40_ and/or Glu_49_; **Figure 6C**). This was supported by a minor loss of PCSK9 activity on the LDLR upon mutation of these residues to Ala (PCSK9-DEE/A), which was not restored under CAP1 overexpression (**Figure 6D**). Furthermore, upon CAP1 overexpression and unlike WT-PCSK9 or PCSK9-3RA, the prodomain mutant PCSK9-DEE/A is inhibitable by P1.40. While PCSK9-Δ33-58 is more active, the combined PCSK9-Δ33-58/3RA is no longer sensitive to CAP1, but is inhibitable by P1.40, like PCSK9-Δ33-58 (**Figure 6D**). Altogether, these results highlight the role of key N-terminal acidic residues in modulating the CAP1 stimulation of the activity of PCSK9.

**Figure 6:**
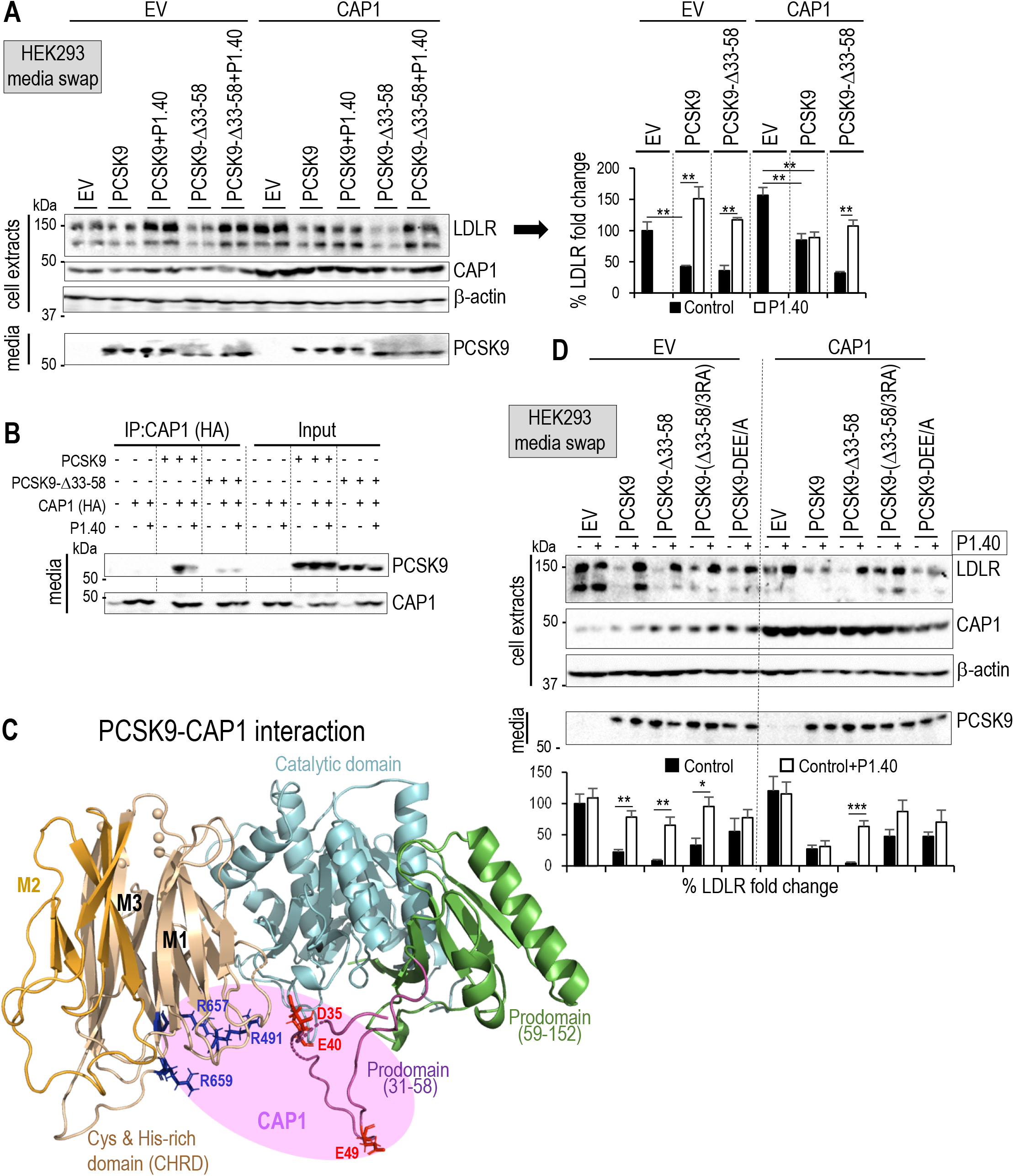
Stimulation by CAP1 and inhibition by P1.40 of PCSK9 activity: validation of CAP1 binding sites in PCSK9 prodomain. (**A**) HepG2-*PCSK9*^*-/-*^ cells were transfected with vectors either empty (EV) or expressing CAP1. After 24h, cells were incubated for 18h with media from HEK293 cells expressing empty vector (EV), PCSK9 (0.3 μg/ml) or PCSK9-Δ33-58 mutant (0.3 μg/ml), in the absence or presence of P1.40 (1 μg/mL). Cell extracts and media were analyzed by Western blot and LDLR signal was quantified. (**B**) EV, PCSK9, PCSK9-Δ33-58, and/or HA-tagged CAP1 were expressed in HepG2 *PCSK9*^−/-^ cells. After 24 h, cells were grown in a serum free medium for 18 h. After immunoprecipitation using HA-coupled agarose beads, immunoprecipitated CAP1-HA and PCSK9 proteins were analyzed by Western blotting and revealed using anti-CAP1 and anti-PCSK9 antibodies respectively (input is 1% of total medium volume used for immunoprecipitation). (**C**) Schematic of CAP1 interaction (pink area) with PCSK9 emphasizing the key role of M1 and M3 (blue), and prodomain (red) specific residues. (**D**) HepG2-*PCSK9*^*-/-*^ cells that expressed an empty vector (EV) or CAP1 for 24h were incubated with the conditioned media from HEK293 cells transfected with an EV or vectors expressing PCSK9 or its mutants (all at 0.3 μg/mL) in the absence or presence of purified P1.40 (1 μg/mL). Cell extracts and media were analyzed by Western blot. Data represent means ± SEM. Representative blots of at least three independent experiments are shown. *P* values (*, *P*<0.05; **, *P*<0.01; ***, *P*<0.001) were obtained by two-way ANOVA coupled to Tukey’s multiple comparisons tests.

Finally, we reported that the natural E670G variant that prevents Ser_668_-phosphorylation at the motif Ser_668_-X-Glu_670_ decreases PCSK9 activity [40]. Interestingly, the E670G variant has an even stronger effect in the absence of endogenous CAP1 (siCAP1; **Supplementary Figure 7E**), suggesting that CAP1 binding to the CHRD may be enhanced by Ser_668_ phosphorylation.

### 3.5 PCSK9 and MHC-I

It was recently reported that PCSK9 targets the mouse and human major histocompatibility complex I (MHC-I) to endosomes/lysosomes for degradation, independently from the LDLR [31]. While PCSK9 binding to the LDLR requires its catalytic domain [11-13], its binding to MHC-I implicates the M2 domain of the CHRD and an exposed **<u>R</u>**-X-**<u>E</u>** motif (Arg_68_ and Glu_70_) in MHC-I receptors, as reported for HLA-C [31]. We modeled the M2 and human HLA-C interaction, leading us to hypothesize that two key residues Glu_567_ and Arg_549_ in M2 may interact with Arg_68_ and Glu_70_ of HLA-C, respectively (**Figure 7A**). To verify whether Arg_549_ and Glu_567_ in M2 may also be important for LDLR degradation, we incubated of HepG2-*PCSK9*^−/-^ cells with PCSK9 or PCSK9-R549A/E567A produced in HEK293 cells. Unexpectedly, the latter double mutation effectively reduced the activity of PCSK9 on the LDLR (**Figure 7B**). This suggests that the M2 domain Arg_549_ and Glu_567_, which likely interact with the **<u>R</u>**-X-**<u>E</u>** motif of HLA-C, are also critical for the PCSK9-induced LDLR degradation.

**Figure 7:**
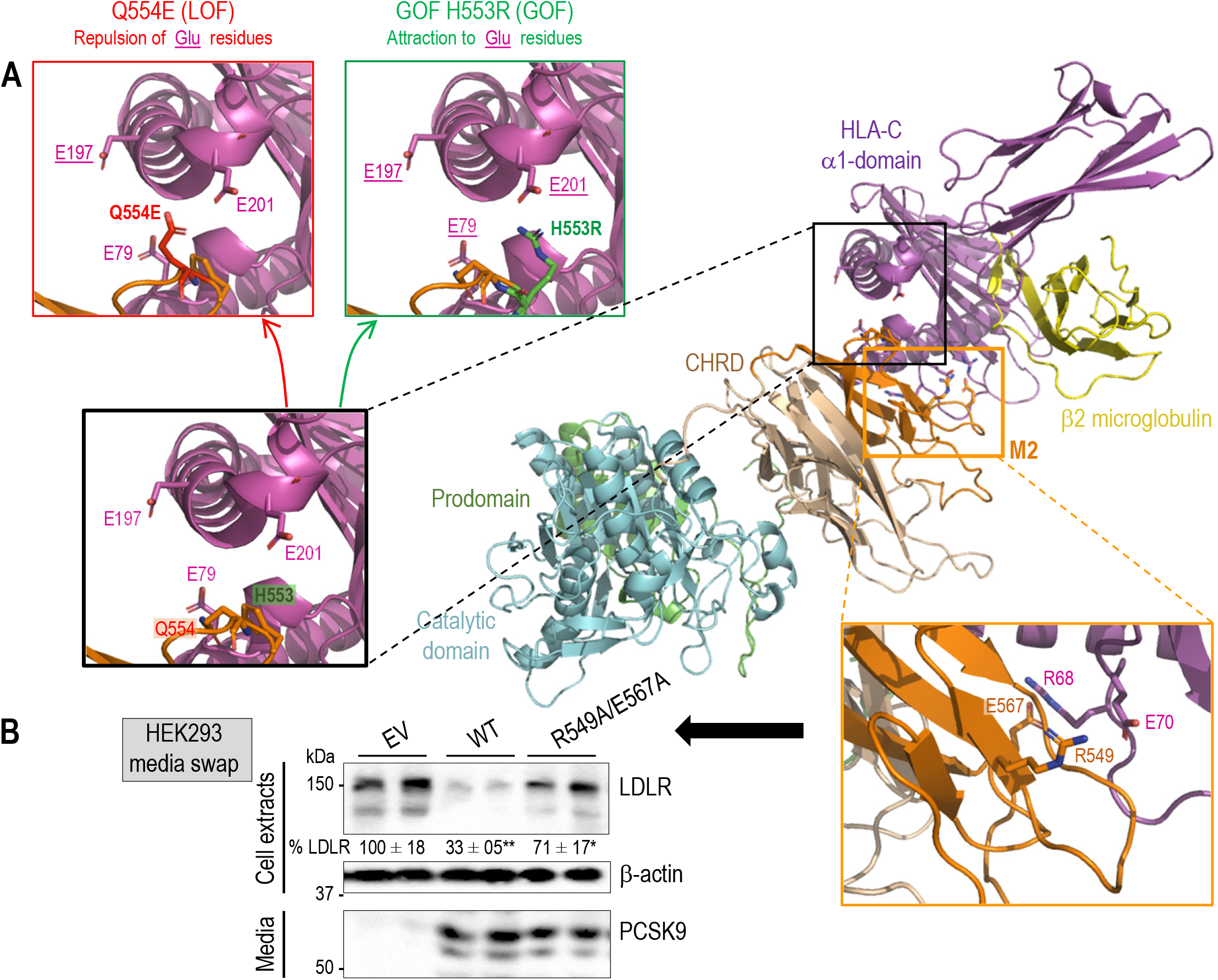
HLA-C: a “protein X” candidate. (**A**) Molecular modeling of the interaction of the M2 subdomain (orange) of PCSK9 with HLA-C (α1 domain; purple). A zoom view shows the proximity of H553 and Q554 to E201, E197 and E79, thereby rationalizing the natural gain-of-function H553R, expected to reinforce the binding of PCSK9 to HLA-C, and loss-of-function Q554E (expected to repulse HLA-C) based on mutagenesis in silico with Pymol. (**B**) Effect of PCSK9-R549A/E567A on the LDLR. HepG2-*PCSK9*^*-/-*^ cells were incubated with conditioned media from HEK293 cells expressing EV, PCSK9 or PCSK9-R549A/E567A mutant (all at 0.3 μg/ml) as indicated. Cell extracts and media were analyzed by Western blot. Data represent means ± SEM. Representative blots of at least three independent experiments are shown. *P* values for LDLR (*, *P*<0.05; **, *P*<0.01) were obtained by two-way ANOVA coupled to Tukey’s multiple comparisons tests.

## 4. DISCUSSION

PCSK9 was shown to bind and enhance the degradation of the LDLR as well as other LDLR family members, such as VLDLR, ApoER2 [55], LRP1 [22], but also the fatty acid transporter and scavenger receptor CD36 [56]. However, the mechanism behind the capacity of PCSK9 to promote the degradation of its target receptors remains obscure. The mechanism behind the PCSK9-induced enhanced extracellular or intracellular [56,57] activities of PCSK9 resulting in the degradation of its target cell surface proteins remains obscure. PCSK9 binds the EGF-A domain of the LDLR *via* its catalytic domain [9-13], but requires its CHRD to escort the LDLR to endosomes/lysosomes for degradation [6,14,15,17-22]. In the latter step, the M2 subdomain of the CHRD is essential [24], suggesting that a membrane-bound “protein X” interacts with M2 to escort the LDLR-PCSK9 complex to degradation compartments [24-26].

Trafficking of the PCSK9-LDLR complex to lysosomes can be disrupted *ex vivo* and *in vivo* by a nanobody P1.40 [27,28], or by a mAb [58] both of which bind the M1 and M3 subdomains, but not M2. This suggests that their binding, while still allowing PCSK9-LDLR interaction, may inhibit the lysosomal sorting of this complex and/or proper exposure of M2, thereby preventing PCSK9 from promoting the degradation of the LDLR [27,28]. This prompted us to analyze the complexes P1.40-PCSK9 (SAXS and BLI) and P1.40-CHRD (X-ray crystallography and BLI). Our data reveal that CHRD-P1.40 interaction relies on two key residues, Asp_480_ in M1 and Arg_105_ in P1.40, as their substitution to Ala eliminated this interaction *in vitro* and in cells. They also show that the flexible hinge [9] between the catalytic domain and the CHRD allows an open conformation of PCSK9, stabilized by P1.40 binding, and a closed conformation largely inaccessible to P1.40 (**Figures 1C, 4E**). This suggested that of these two interconvertible structures [9], the closed conformation best promotes LDLR degradation.

The search for natural CHRD-binding proteins identified two cytosolic, but secretable proteins: annexin A2 [59] and CAP1 [30]. Annexin A2 binds Gln_554_ in the M2 domain of the CHRD [59] and inhibits the activity of PCSK9 on the LDLR in non-liver tissues [60]. The structural similarity of the CHRD to resistin [61] led to the identification of the resistin-binding CAP1 [29] as a potential CHRD-binding protein that enhanced PCSK9 activity on the LDLR [30]. Befittingly, CAP1 mRNA levels represent 50% and 150% of PCSK9 ones in mouse hepatocytes (single cell RNA-seq; https://tabula-muris.ds.czbiohub.org/) or liver (https://www.ncbi.nlm.nih.gov/geo/query/acc.cgi?acc=GSE205008) [62], respectively. It was proposed that the primarily cytosolic CAP1 could somehow associate with the extracellular surface of hepatocytes by an unspecified membrane-flipping mechanism and hence interacts with extracellular PCSK9 at the cell surface [30].

In this study, we showed for the first time that CAP1 is well secreted from hepatic cell lines (**Figures 2A, 3A**), possibly by an unconventional secretory pathway [63] that, like for Annexin-A2, might implicate caveoli [64]. Indeed, CAP1 was reported to circulate in the plasma of metastatic cancer [65] patients and in mouse models of breast cancer [66]. We suggest that the ability of CAP1 to enhance PCSK9 activity relies on its secreted form that would bind PCSK9 extracellularly. Furthermore, it was suggested that in the presence of CAP1 the PCSK9-LDLR degradation in HepG2 cells requires endocytosis in caveolin-coated vesicles [30]. Our data differ. In agreement with our previous results in HepG2 and other cell lines [14,51], we show that the PCSK9-LDLR complex is endocytosed into clathrin-coated vesicles and is not caveolin-dependent (**Figure 2B**,**D**). Caveoli may rather be necessary for the secretion of the cytosolic CAP1 [63,64].

The present data show that extracellular CAP1 binds the M1 and M3 subdomains of PCSK9, like P1.40, but at distinct sites. We identified three arginines in the M1 (Arg_491_) and M3 (Arg_657_ and Arg_659_) domains, as well as three acidic residues (Asp_35_, Glu_40_ and Glu_49_) in the prodomain [53,54], as relevant CAP1 binding sites. The prodomain site seems to be the most important, since the activity of PCSK9-3RA mutant is rescued by CAP1, whereas PCSK9-DEE/A is no longer sensitive to CAP1 (**Figure 6D**). We hypothesized that, even though both CAP1 and P1.40 can each bind PCSK9 in the open conformation **(Figure 4E**), CAP1 likely stabilizes the closed conformation of PCSK9 (**Figure 4E**), thereby preventing access of P1.40 to the key Asp_480_ of PCSK9. Under these conditions, CAP1 can abrogate the P1.40-mediated inhibition of the PCSK9 activity on the LDLR (**Figure 5**). Our SAXS studies further revealed that PCSK9 predominantly adopts a “closed” conformation at neutral pH. Altogether our data indicate that CAP1 binding to PCSK9 favours M2 domain exposure to “protein X” that likely drags the LDLR-PCSK9-CAP1 complex from early endosomes to lysosomes for degradation. This is supported by the absence of CAP1 stimulation of the double mutant PCSK9-Δ33-58 + PCSK9-3RA that lacks CAP1 binding sites (**Figure 6D**; **Supplementary Figure 7C**). In the segment 33-58 of the prodomain of PCSK9, there are multiple acidic residues, a sulphated Tyr_38_ and a phosphorylated Ser_47_ [54]. However, their single or collective contributions to the modulation of PCSK9 activity remain to be defined [67]. CAP1 mRNA silencing significantly reduced PCSK9 activity towards LDLR (**Figures 2, 3**), but did not abrogate it as originally suggested [30], revealing that CAP1 is an important but non-essential facilitator in this process. Since CAP1 lacks a cytosolic tail to potentially participate in the escort of the LDLR-PCSK9-CAP1 complex to lysosomes, it is not likely to be the sole component of the elusive “protein X” [30].

PCSK9 was recently implicated in MHC-I degradation *via* the interaction of an **R**-X-**E** motif with M2. Our molecular modeling suggests that Arg_68_-X-Glu_70_ of the MHC-I family member HLA-C [26] may interact with M2 Glu_567_ and Arg_549_, respectively (**Figure 7A**). Remarkably, mutation to Ala of these two residues (E567A+R549A) prevented LDLR degradation (**Figure 7B**). Thus, we propose that the “protein X” is likely HLA-C in hepatocytes. This hypothesis is reinforced by our modeling that rationalizes the effect of two neighboring natural gain-of-function (H553R) and loss-of-function (Q554E) variants of PCSK9 [59,68], since they would respectively attract and repulse Glu_79_, Glu_197_ and Glu_201_ in HLA-C (**Figure 7A**). The fact that the LDLR is resistant to PCSK9-induced degradation in CHO cells [14] that lack HLA-C expression [69] also reinforces our hypothesis. Notably, the mouse HLA-C (H2-K1) and its chaperone β2-microglobulin (β2M) are more expressed than PCSK9 in hepatocytes (by 45- and 62-fold, respectively) and in the liver (by 30- and 20-fold, respectively). Because of these large excesses, the identification of HLA-C and/or other MHC-I members as strong candidates for the long-sought “protein X” cannot be unambiguously confirmed by siRNA silencing and will rather require a more efficient CRISPR-Cas gene-inactivation approach in hepatocytes.

## 5. CONCLUSIONS AND FUTURE PERSPECTIVES

The discovery of PCSK9 in 2003 [2] and its implication in the regulation of LDLc [3] was a game changer in the development of more efficient treatments of hypercholesterolemia and associated cardiovascular diseases [26,70,71]. Current treatments combining statins with either subcutaneously injected monoclonal antibodies or siRNA directed to the liver, result in 50-60% further reduction in circulating LDLc above that obtained with statins alone. Whether such an approach could be applied to the treatment of cardiovascular diseases as well as other pathologies such as immune regulation in cancer/metastasis [72,73] begs future investigations.

Our present structural and functional data suggest that the sorting of the PCSK9-LDLR to endosomes/lysosomes for degradation is a complex process that implicates specific residues within an exposed M2 domain of the CHRD linking the complex to an uncharacterized “protein X”. The present data extend our understanding of sorting mechanisms linking directional changes in the PCSK9-induced degradation of specific cell surface receptors. This process is positively modulated by CAP1 that binds the CHRD M1 and M3 domain as well as specific acidic residues within the PCSK9 prodomain, thereby twisting PCSK9 into a closed conformation that best presents the M2 domain for subcellular sorting to endosomes/lysosomes. The similarity of this regulatory process to the one used in the sorting of PCSK9-MHC-I complex for lysosomal degradation suggests a common regulatory mechanism induced by extracellular PCSK9. This mechanism is likely distinct from the intracellular pathway of PCSK9-LDLR degradation, as the latter is not dependent on the M2 domain [24]. Our study opens the way to the design of potent and specific M2 domain binding proteins or small molecules that would inhibit the sorting of PCSK9 to lysosomes and hence its ability to escort specific surface receptors to degradation compartments.

## Supporting information

Matgerials and Methods and Supplmental Figures and Tables

## Abbreviations

β2M: β2-microglobulin
CAP1: cyclase associated protein 1
CHC: clathrin heavy chain
CHRD: cysteine histidine rich domain
CRISPR: clustered regularly interspaced short palindromic repeats CTL control
EV: empty vector
FACS: fluorescence-activated cell sorting
HLA-C: human leukocyte antigen-C
LDLR: low density lipoprotein receptor
MHC-I: major histocompatibility class-I
PCSK9: proprotein convertase substilisin kexin 9
rPCSK9: recombinant PCSK9

## DATA STATEMENT

Data will be made available on request.

## AUTHOR CONTRIBUTIONS

Conceptualization, C.F.G. and N.G.S.; Investigation, C.F.G., A.B.D.O., L.C., E.G., S.M., A.E., M.L.D., E.C.L., H.N., O.P.R., A.T., D.S-R., and P.L.; Formal Analysis and Visualization, C.F.G., A.B.D.O., A.P., V.D., and N.G.S., Original Draft, N.G.S., C.F.G., A.B.D.O.; Review & Editing, N.G.S., A.P., C.G.F.; Funding Acquisition and Project Administration, N.G.S.; Supervision, N.G.S. N.G.S. takes primary responsibility for the data described in this manuscript.

## ACKNOWLEDGEMENTS

This work was supported in part by a CIHR Foundation grant (NGS: # 148363), a Canada Research Chairs in Precursor Proteolysis (NGS: # 950-231335) and a Leducq Foundation grant (NGS: # 13 CVD 03). Sepideh Mikaeeli was supported by an FRQS fellowship. We acknowledge SOLEIL for provision of synchrotron radiation facilities, and we would like to thank the staff of beamlines SWING, PROXIMA-1, and PROXIMA-2 for assistance in setting up our experiments. The authors would also like to acknowledge the expert secretarial help of Habiba Oueslati in the preparation of this manuscript.

## CONFLICT OF INTEREST

None declared.

## APPENDIX A. SUPPLEMENTARY DATA

Supplementary data to this article can be found online at https://

### Online Methods

For a detailed description of all methods, see the Supplementary material online, Experimental Procedures.

### Accession codes

The atomic coordinates have been deposited at the Protein Data Bank, with accession codes 7ANQ.

## Graphical Abstract

Model representing the interactions of PCSK9 with LDLR (EGF-A domain), P1.40 (M1, M3), CAP1 (M1, M3 and prodomain) and “Protein X” (M2 domain).

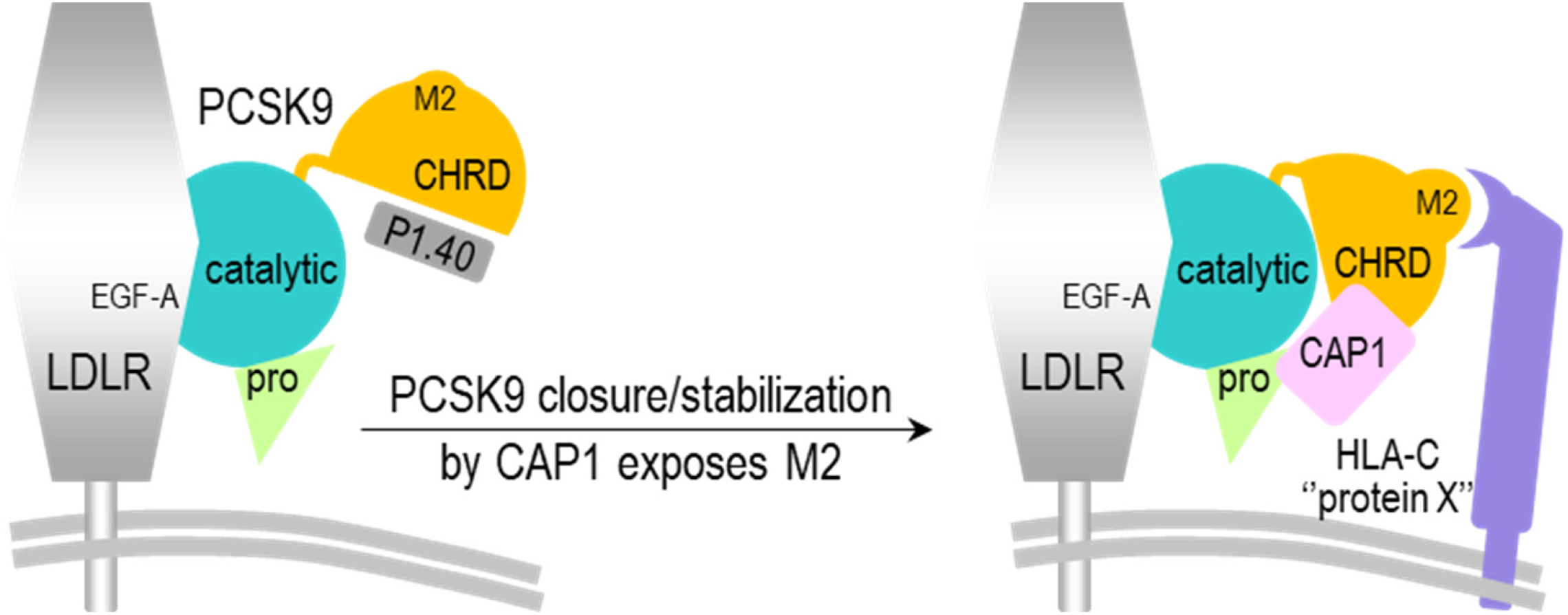

## Supplementary Figures

**Supplementary Figure 1:**
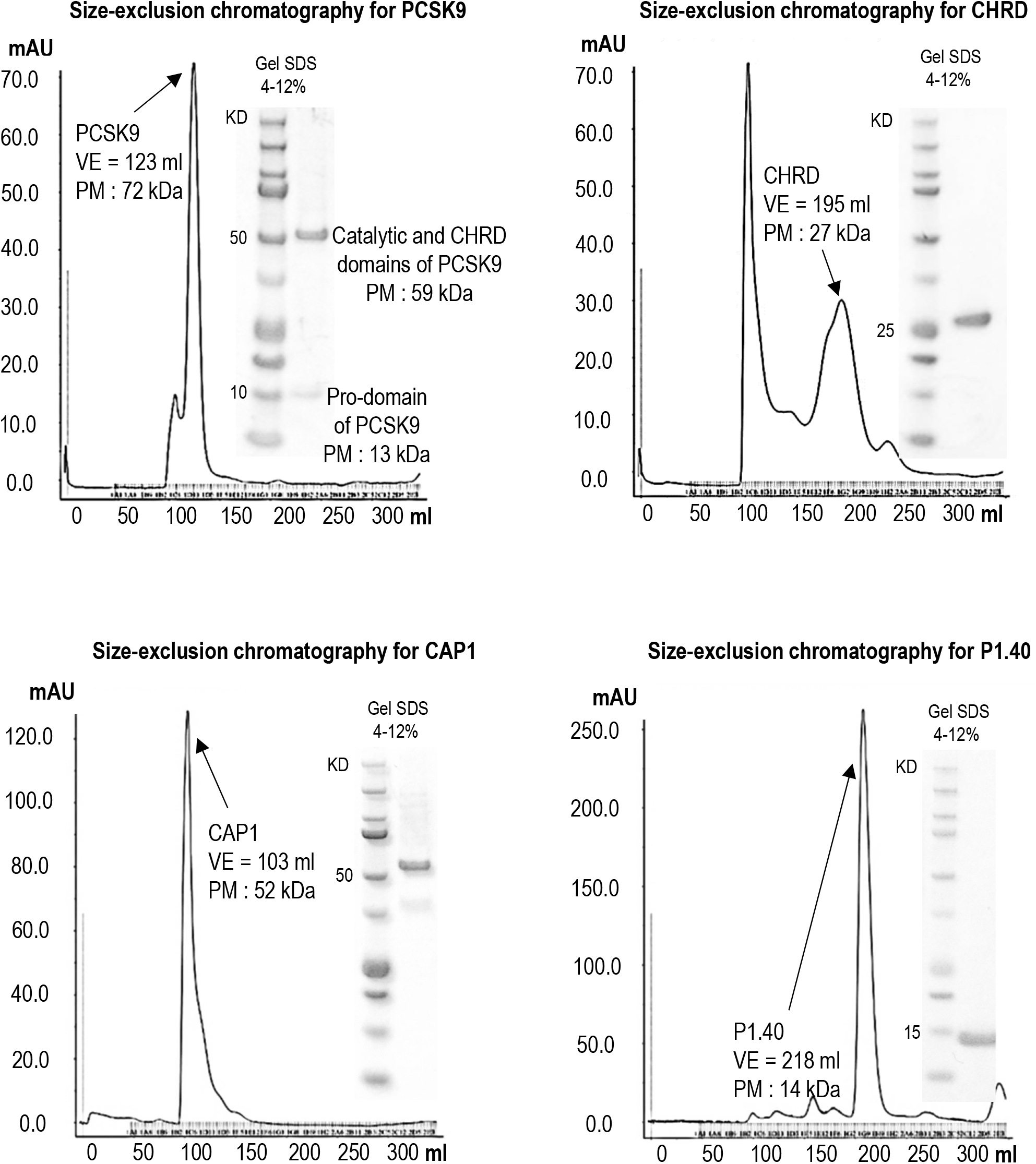
PCSK9, CHRD, CAP1 and nb P1.40 purified from Size-exclusion chromatography (SEC) Sephacryl ® S-100 HR GE Healthcare in buffer 50 mM Tris HCl pH 8, 150 mM NaCl. and loaded on 4-12% SDS PAGE.

**Supplementary Figure 2:**
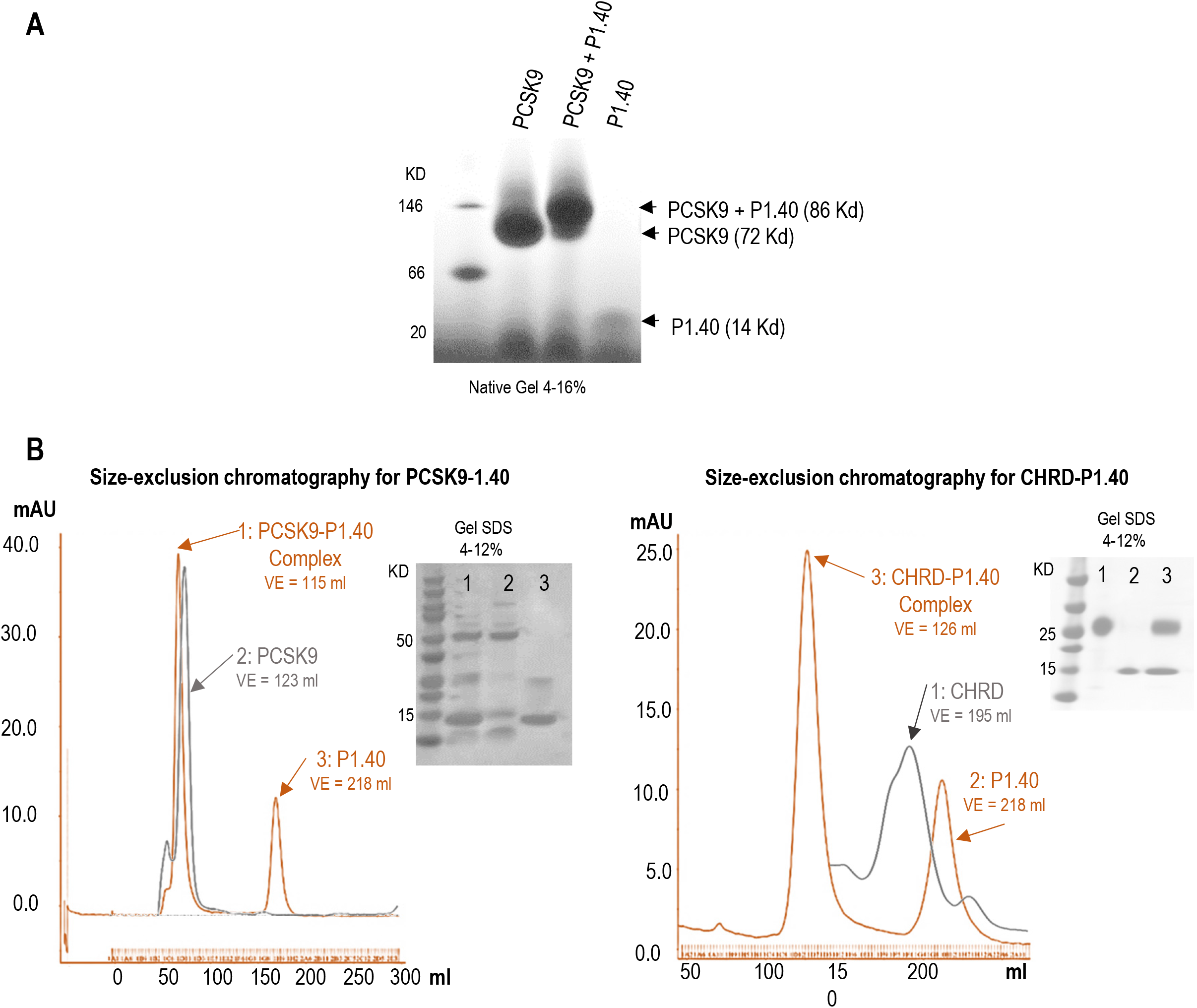
Characterization of the PCSK9-P1.40 complex. **(A)** NativePage™ 4-16% Bis-Tris Ge.(B) Purification of PCSK9-P1.40 and CHRD-P1.40 by Size-exclusion chromatography (SEC) Sephacryl ® S-100 HR GE Healthcare in buffer 50 mM Tris HCl pH 8, 150 mM NaCl and loaded on 4-12% SDS PAGE.

**Supplementary Figure 3:**
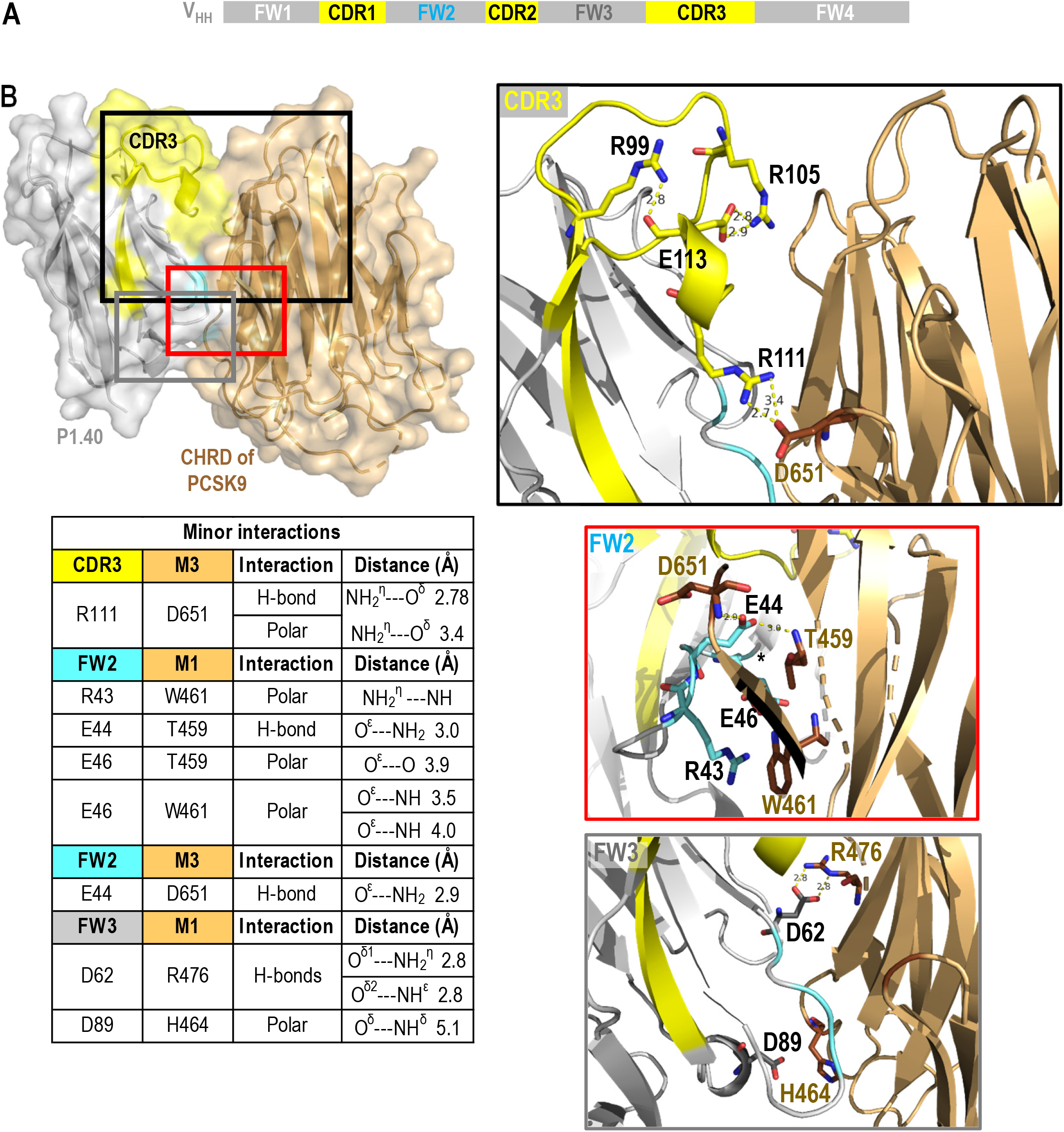
Analysis of P1.40-CHRD interactions in the crystal complex PDB 7ANQ. (**A**) Structure of the variable heavy chain-only (V_HH_) antibody P1.40 raised in llama [8]. The antigen binding site is constituted of three complementary-determining regions (CDRs) framed with framework regions (FWs) implicated in the core structure of the antibody. (**B**) Surface representation of the P1.40 (grey)-CHRD (light brown) crystal complex and modeling of additional interactions established between P1.40 domains CDR3 (yellow), FW2 (cyan) and FW3 (dark grey) and the CHRD of PCSK9, which is composed of M1, M2 and M3 subdomains. **CDR3** (aa 105-122): between Arg_111_ and M3 Asp_651_. **FW2**: between Arg_43_, Glu_44_, Glu_46_ and M1 Thr_459_, Trp_461_ or M3 Asp_651_. **FW3**: between Asp_62_, Asp_89_ and M1 His_464_, Arg_476_.

**Supplementary Figure 4:**
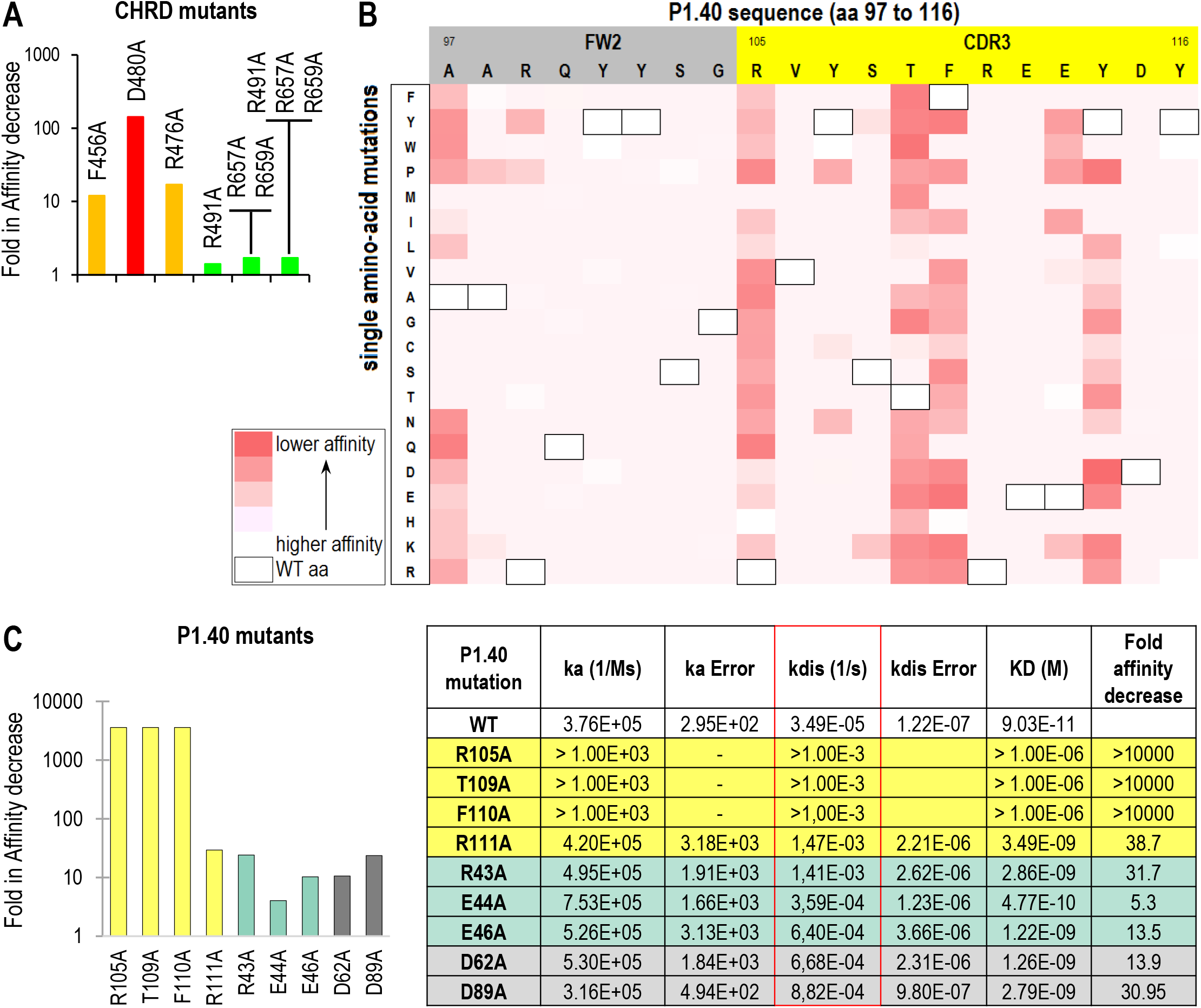
Identification of the key interacting residues in the P1.40-CHRD complex. (A) Decreased affinity was measured by yeast cell-surface display of P1.40 and CHRD binding after Ala replacement of selected CHRD residues. (**B**) A library containing every possible amino acid change at each position of a selected region of P1.40 (Deep Mutational Scanning) was generated by PCR and tested for the loss of capacity of each P1.40 mutant expressed at the yeast cell surface to bind the CHRD (negative sorting). White color indicates that P1.40-CHRD binding was unchanged while the darkest red indicate a loss of binding. (**C**) Impact of targeted mutations on biotinylated P1.40 affinity for the CHRD are shown. Corresponding kinetic constants were analyzed by BLI using a streptavidin sensor.

**Supplementary Figure 5:**
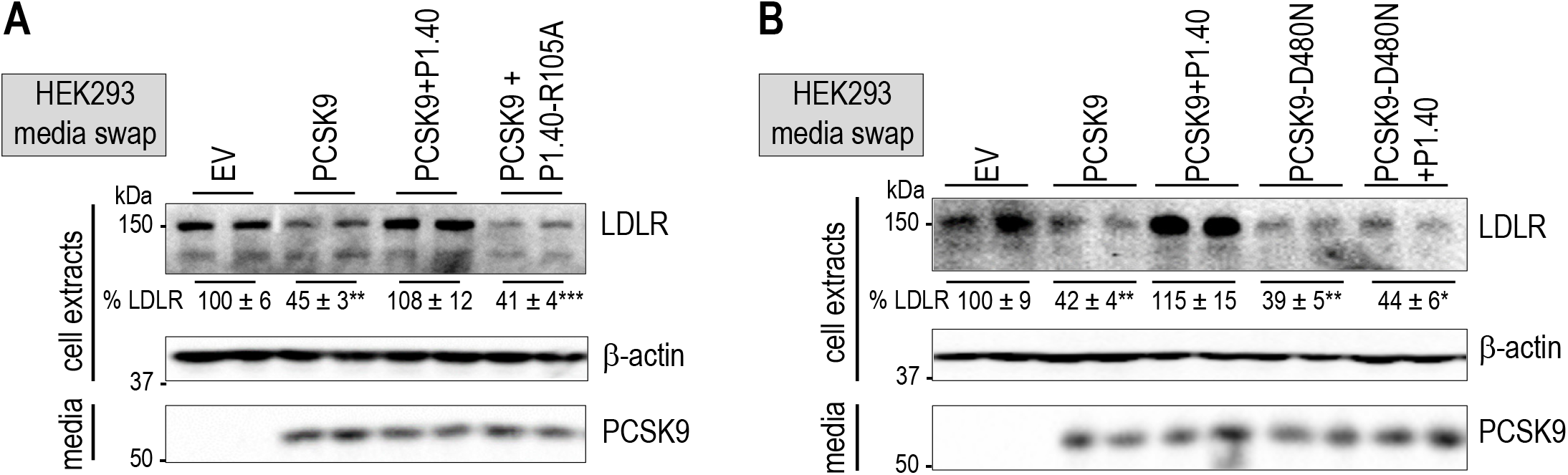
Cell validation of the key interaction between the Arg_105_ in P1.40 and Asp_480_ for the inhibitory effect of P1.40. (**A**) HepG2-*PCSK9*^−/-^cells were incubated with conditioned media from HEK293 cells expressing an empty vector (EV) or PCSK9-V5 (0.3 μg/ml), in the absence or presence of purified P1.40 or P1.40-R105A mutant (1 μg/ml). LDLR protein levels were normalized to β-actin and set to 100 for EV. (**B**) The same conditions as in (A) were used, except that PCSK9 or PCSK9-D480N were expressed and used at 0.3 μg/ml. Means ± SEM for three (A) or two (B) independent experiments. *P* values for LDLR (*, *P*<0.05; **, *P*<0.01; ***, *P*<0.001) were obtained by two-way ANOVA coupled to Tukey’s multiple comparisons tests.

**Supplementary Figure 6:**
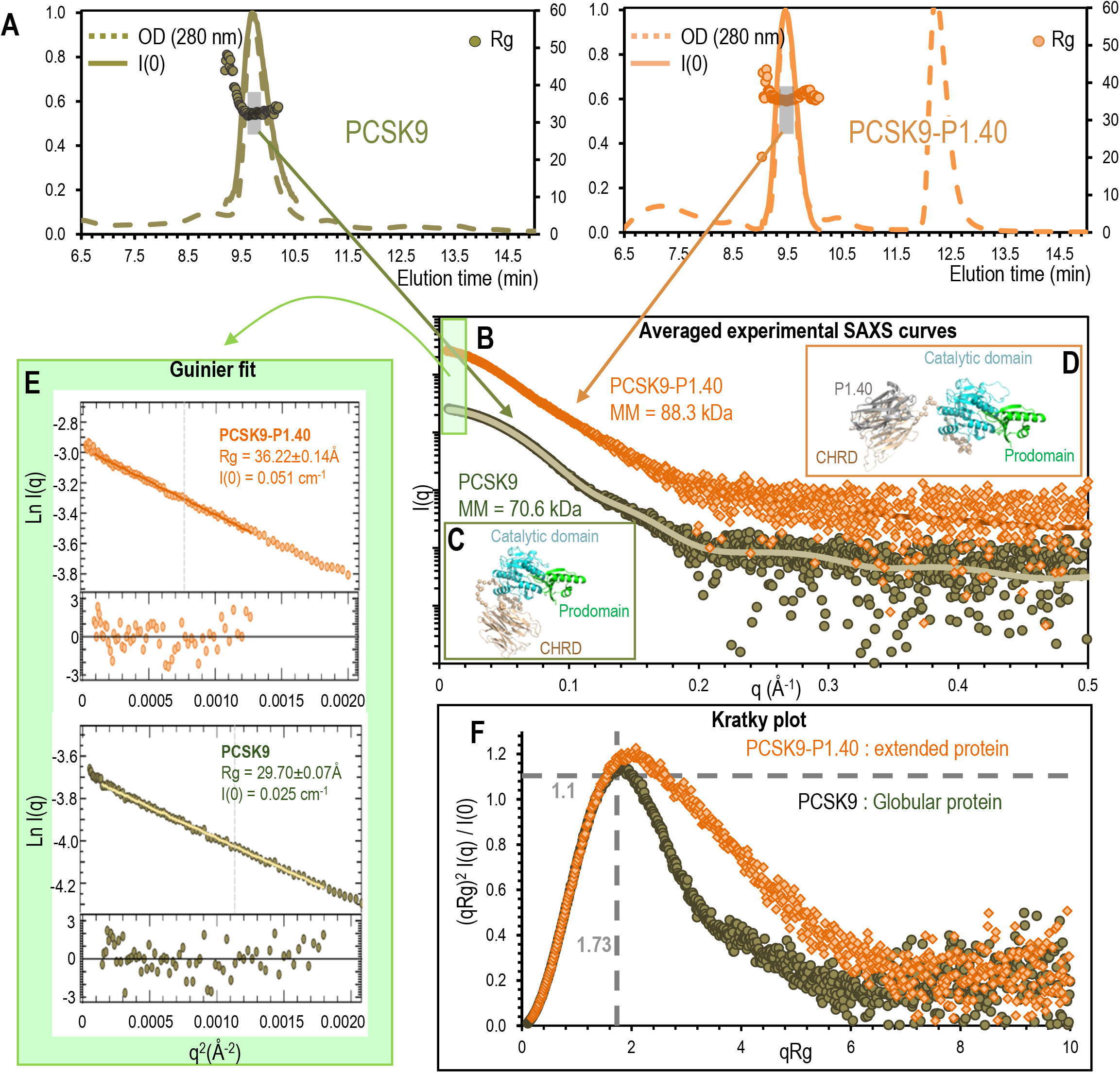
Size exclusion chromatography and SAXS data. (**A**) UV (280 nm), I(0) (vector intensity; solid) and Rg (radius of gyration; dots) elution profiles of the protein PCSK9 alone (green) and in complex with P1.40 (orange). Normalized UV and I(0) values are shown on the left y-axis and Rg values in Å on the right one. OD_280nm_ profiles indicates that PCSK9 is monomeric in solution, peaking at 9.8 min, and that the PCSK9/P1.40 complex is stable, does not dissociate and peaks at 9.5 min. The second peak at 12.3 min corresponds to excess monomeric P1.40. Selected fractions for SAXS curves averaging (9.5 to 10 min and 9.3 to 9.7 min for PCSK9 alone and in complex, respectively) are shown in grey. (**B**) Experimental SAXS curves were obtained by averaging and defined scattering vectors in Å^−1^ (x-axis) and their intensities (y-axis). The CRYSOL [9] results fitting (green and orange lines) with the best DADIMODO [10] models for PCSK9 alone (**C**) and in complex with P1.40 (**D**) gave a χ^2^ of 1.15 and 0.97 respectively. (**E**) The light green rectangle in (B) corresponds to the selected area for the Guinier fit [11] for q closed to 0. The Guinier law defines the radius of gyration Rg and I(0). (**F**) In the normalized Kratky plot [12,13] making it possible to quickly assess the globular nature and the flexibility, the dashed cross represents the theoretical value for the maxima of a globular protein (peaks at 1.1 for qRg of 1.73). The experimental curves (dot) show that the global shape of the complex with a maximum peak at 1.2 (orange symbols) is more extended compared to the global shape of the protein PCSK9 alone (green symbols) with a maximum at 1.16. The estimation of molecular weight of PCSK9 alone (70.6 kDa) and of the PCSK9/P1.40 complex (88.3 kDa) by SAXS using the Bayesian estimation from ATLAS [14] indicates that a single protein P1.40 (14.4 kDa) binds to PCSK9 (72.5 kDa) in solution.

**Supplementary Figure 7:**
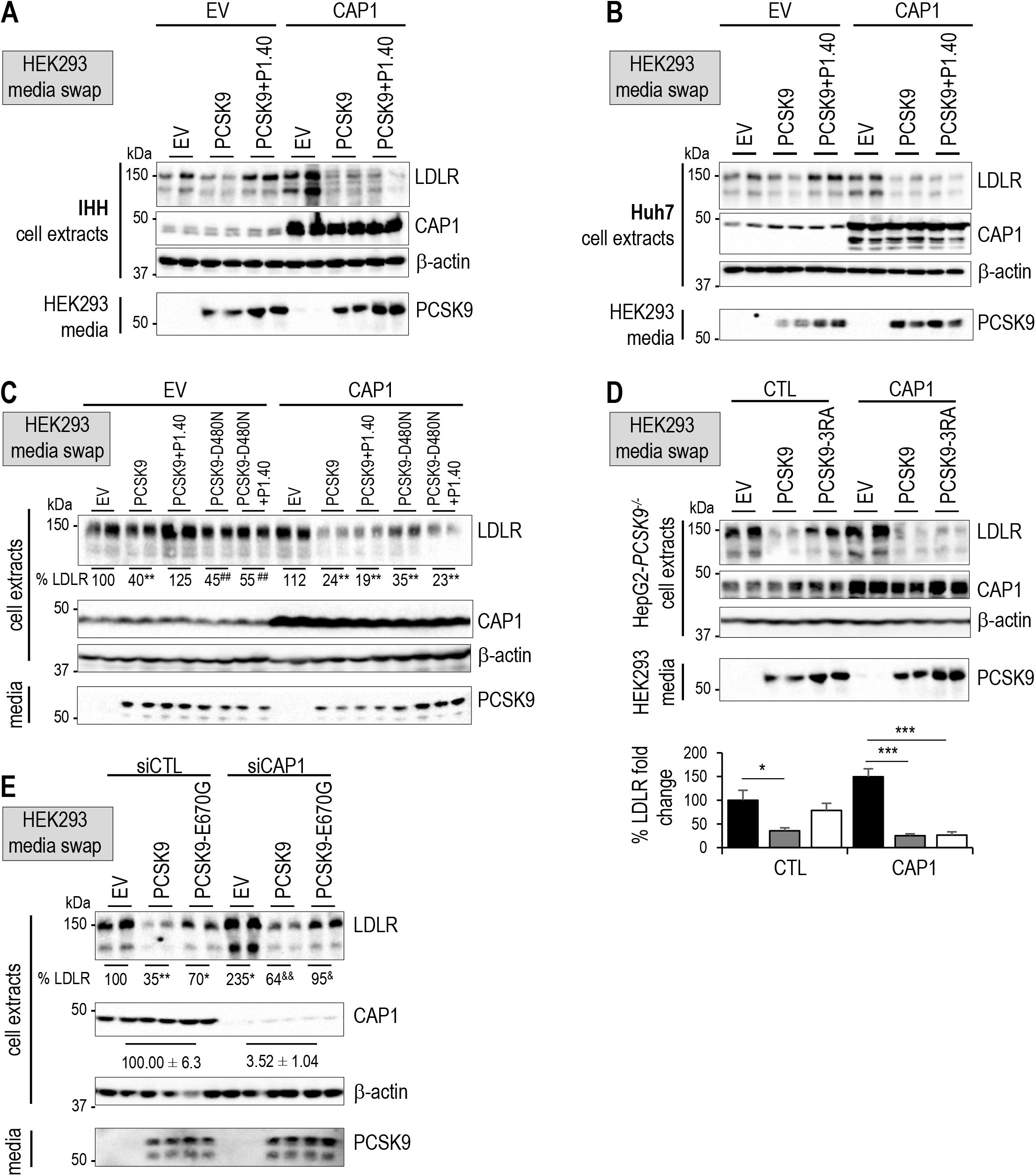
Regulation of PCSK9 activity by CAP1 and P1.40. IHH (**A**) or Huh7 (**B**) cells were transfected for 24 h with an empty-(EV) or CAP1-expressing vector and then exposed for 18 h to conditioned media from HK293 cells, as in Figure 5A. (**C**) HepG2-*PCSK9*^*-/-*^ cells expressing an empty vector (EV) or CAP1 (24 h) were treated for 18 h with conditioned media from HEK293 cells expressing an empty vector (EV) or incubated with PCSK9 or PCSK9-D480N (0.3 μg/mL) in combination with P1.40 (1 μg/ml). (**D**) HepG2-*PCSK9*^*-/-*^ cells were treated under the same conditions as in (C) with PCSK9 or PCSK9-3RA. (**E**) HepG2-*PCSK9*^−/-^ cells were transfected with siRNA constructs including siCTL or siCAP1 and incubated with conditioned media from HEK293 cells expressing an empty vector (EV), PCSK9 or PCSK9-E670G mutant (0.3 μg/ml) for 18 h. Cell extracts and media were analyzed by Western blot. Representative blots of three (A-D) and two (E) independent experiments are shown. Data represent means ± SEM (*, *P*<0.05; ***, *P*<0.001; ^*^, *P*<0.05; ^*^, *P*<0.01) obtained by two-way ANOVA coupled to Tukey’s multiple comparisons tests.

## Notes

### Competing Interest Statement

The authors have declared no competing interest.

