## Supplementary material for "Molecular interactions of PCSK9 with an inhibitory nanobody, CAP1 and HLA-C: functional regulation of LDLR levels": Matgerials and Methods and Supplmental Figures and Tables

**Supplementary Table 1**. Primers used for various PCSK9 mutants.

| Construct | Restriction enzyme set | Primers (5' – 3') |
|---|---|---|
| PCSK9-DEE/A (D35A-E40A-E49E) | *SacII/SalI* | *5'- GATGTAGTCGACATGGGGCAACTTCAAGGCCA GCTCCAGCAGGTCG-3'* *5'-GGGTCCCGCGGGCGCCCGTGCGCAGGAGGA CGAGGCCGGCGACTACGAGGCGCTGGTGCTAGCCTTG CGTTCCGAGGCGGACGGCCTGGCCGAAG -3'* |
| PCSK9-CHRD (aa 449-693) | *AgeI/SacI* | *5'*-AGGGTG CCAGCATGCGCAG-*3'* *5'-GGCTTACCGGTCTG GGGTAGCAG GCAG-3'* |
| PCSK9-D480N | *AgeI/SacI* | *5'-GCCCCAAATGAGGAGCTGCTG-3'* *5'-CTCCTCATT TGGGGCGCAGC-3'* |
| PCSK9-R491A | *AgeI/SacI* | *5'-CTTCCCACTCGCGGAGAAAC-3',* *5'-GTTTCTCCGCGAGTGGGAAG-3'* |
| PCSK9-R549A/E567A | *AgeI/SacI* | *5'-GTGCTGACGTCCGCGCTCGCGACTACACACG-3'* *5'-CGTGTGTAGTCGCGAGCGCGGACGTCAGCAC-3'* |
| PCSK9-R657A/R659A | *AgeI/SacI* | *5'-TGCTGACGTCCGCGCTCGCGACTACACACG-3'* *5'-CGTGTGTAGTCGCGAGCGCG GACGTCAGCA-3'* |
| PCSK9-3RA R491A/R657A/R659A | *AgeI/PshAI* | Ligation of resulting fragments from PCSK9-R491A and PCSK9FL-R657A/R659A constructs |
| PCSK9-Δ33-58/3RA | *AgeI/SacI* | Ligation of resulting fragments from PCSK9-Δ33-58 and PCSK9-R491A-R657A-R659A constructs |

**Supplementary Table 2**. Crystallographic data and refinement statistics.

| ***Structure*** | **CHRD/P1.40 complex** |
|---|---|
| ***PDB code*** | **PDB 7ANQ** |
| ***Data Collection Source*** | SOLEIL PROXIMA-1 |
| Space group | $P4_12_12$ |
| ***Unit Cell*** | |
| $a = b$, $c$ (Å) | 52.3, 265.2 |
| $\alpha = \beta = \gamma$ (°) | 90 |
| Wavelength (Å) | 0.978570 |
| Resolution (Å) | 2.196 (2.196-2.33) |
| No. of reflections | 407293 |
| No. of unique reflections | 19768 |
| Completeness (%) | 100 (99.8) |
| $I/\sigma(I)$ | 12.04 (2.05) |
| $CC_{1/2}$ | 0.997 (0.683) |
| $R_{merge}$ (%) | 24.6 (172.9) |
| Multplicity | 20.60 (20.80) |
| ***Refinement*** | |
| Resolution (Å) | 2.20 |
| $R_{work}$ | 0.213 |
| $R_{free}$ | 0.256 |
| R.m.s. bond length deviation (Å) | 0.008 |
| R.m.s. bond angle deviation (°) | 1.09 |
| Ramachandra outliers (%) | 0.00 |

All data set were processed using XDS [1,2] with the xdsme script (*Legrand, P. XDSME: XDS Made Easier-GitHub Repository, 2017; https://github.com/legrandp/xdsme and https://doi.org/10.5281/zenodo.837885*), before being scaled together with XSCALE and run through XDSME/XDSCONV to generate MTZ files. The structure was solved by molecular replacement using Phaser [3], starting from PDB 3H42 and 4EIZ as models, followed by refinement using REFMAC5 [4]. The electron density maps were interpreted using COOT [5]. The structures were subjected to at least three cycles of rebuilding and refinement performed using with REFMAC5 [4], PHENIX.refine [6] and BUSTER [7]. The figures were made with PyMOL (*DeLano, W. L. The PyMOL Molecular Graphics System (Schrödinger, L., New York Editor, 2010*).

**Supplementary Table 3**. SAXS analysis of PCSK9 alone or in complex with P1.40.

<table>
<tr><th colspan="4">SAXS data collection parameters</th></tr>
<tr><td colspan="2">Source, instrument and description or reference</td><td colspan="2">SOLEIL/SWING, U20 in-vacuum undulator</td></tr>
<tr><td colspan="2">Wavelength (Å)</td><td colspan="2">1.033</td></tr>
<tr><td colspan="2">Beam geometry (size, sample-to-detector distance)</td><td colspan="2">400 x 200μm$^2$</td></tr>
<tr><td colspan="2">$q$-measurement range (Å$^{-1}$)</td><td colspan="2">0.0073-0.58<br>($q = 4\pi\sin\theta.\lambda^{-1}$, where 2θ is the scattering angle)</td></tr>
<tr><td colspan="2">SAXS Detector</td><td colspan="2">EigerX 4M in vacuum</td></tr>
<tr><td colspan="2">Absolute scaling method</td><td colspan="2">Water</td></tr>
<tr><td colspan="2">Basis for normalization to constant counts</td><td colspan="2">Active beamstop : diamond based diode</td></tr>
<tr><td colspan="2">Exposure time,<br>Number of exposures</td><td colspan="2">0.99s (0.01s dead time).<br>180 + 720 (buffer+sample)</td></tr>
<tr><td colspan="2">Sample configuration including path length and flow rate</td><td colspan="2">Flowing capillary – 1.5mm of Internal Diameter<br>0.3ml/min</td></tr>
<tr><td colspan="2">Sample temperature (ºC)</td><td colspan="2">15°C</td></tr>
<tr><th colspan="4">Software employed for SAS data reduction, analysis and interpretation</th></tr>
<tr><td colspan="2">SAXS data reduction to sample–solvent scattering, and extrapolation</td><td colspan="2">Foxtrot (3.5.2)<br>US-SoMo (9.9-3106)</td></tr>
<tr><td colspan="2">SAXS data analysis</td><td colspan="2">Atsas suite (3.0.1)</td></tr>
<tr><td colspan="2">Molecular Modelling</td><td colspan="2">MODELLER (9.18) & DADIMODO</td></tr>
<tr><th colspan="4">Structural parameters</th></tr>
<tr><td colspan="2">Sample</td><td>PCSK9</td><td>PCSK9-P1-40</td></tr>
<tr><td rowspan="3">Guinier Law</td><td>Rg (Å)</td><td>29.70 (+- 0.07)</td><td>36.22 (+-0.14)</td></tr>
<tr><td>I(0) (cm$^{-1}$)</td><td>0.025 (+-0.000)</td><td>0.051 (+-0.000)</td></tr>
<tr><td>$q_{max}Rg$</td><td>1.26</td><td>1.29</td></tr>
<tr><td rowspan="5">Distance Distribution Function P(r)</td><td>I(0) (cm$^{-1}$)</td><td>0.025 (+- 0.000)</td><td>0.051 (+- 0.001)</td></tr>
<tr><td>Rg (Å)</td><td>29.76 (+- 0.06)</td><td>37.17 (+- 0.12)</td></tr>
<tr><td>Dmax (Å)</td><td>103</td><td>129</td></tr>
<tr><td>$q_{range}$ (Å$^{-1}$)</td><td>0.004-0.269</td><td>0.009-0.221</td></tr>
<tr><td>$\chi 2$</td><td>0.90</td><td>0.87</td></tr>
<tr><td rowspan="2">Molecular Weight (kDa)</td><td>From Bayesian estimation</td><td>70.6</td><td>88.3</td></tr>
<tr><td>From primary sequence</td><td>72.5</td><td>86.9 (72.5+14.4)</td></tr>
<tr><td rowspan="3">MODELLER<br><br>DADIMODO<br>(15 generated models)</td><td>Starting crystal structures: $\chi 2$</td><td>2P4E (closed) : 1.46<br>2P4E (open) : 75.96</td><td>2P4E + 7ANQ: 3.19</td></tr>
<tr><td>Flexible residues</td><td>30-60, 422-454 & 682-703</td><td>30-60, 422-454 & 682-703</td></tr>
<tr><td>$\chi 2$ range</td><td>1.15-1.26</td><td>0.97-1.05</td></tr>
</table>

Supplementary Figures.

**Supplementary Figure 1: Size-exclusion chromatography.** PCSK9, CHRD, CAP1 and nb P1.40 purified from Size-exclusion chromatography (SEC) Sephacryl ® S-100 HR GE Healthcare in buffer 50 mM Tris HCl pH 8, 150 mM NaCl. and loaded on 4-12% SDS PAGE.

### Experimental Procedures.

**Quantitative RT-PCR (QPCR)**

QPCR of hLDLR mRNA was carried out as previously described [15] using the following primers: *5'*-CCATATGCATCCCCAGTCTT-*3'*, *5'*-AATCCATCTTGTTCAATG GCCGATC-*3'*.

**Cell binding assays for determining the affinity of 6xHis-CHRD variants for P1.40-HA**

*Yeast display pNT P1.40 vector construction:* The pNT P1.40 plasmid allowing the functional display on the surface of *S. cerevisiae* cells is derived from pCT L.7.5.1 (Addgene #42900) which include the Aga2p signal peptide, the P1.40 sequence flanked by NheI and NotI restriction sites, an HA tag for detection and Aga2p sequence for tethering to the yeast cell wall.

*Yeast transformation:* Preparation of competent yeast cells EBY100 /ATCC® MYA-4941™ (MATa AGA1::GAL1-AGA1::URA3 ura3-52 trp1 leu2-delta200 his3-delta200 pep4::HIS3 prbd1.6R can1 GAL) was performed according Benatuil *et al.* [16] and Suga *et al.* [17]. Then, 100 ml of EBY100 electro-competent cells were mixed with 1-2 µg plasmid DNA, transferred to a pre-chilled electroporation cuvette (Biorad, 165–2086) and pulsed at 2.5 kV, 25 mF (Biorad Gene Pulser Xcell). Yeast cells were then diluted in 1 ml of sorbitol (1 M), and 200 ml were streaked on SD-CAA agar plates (6.7 g/l yeast nitrogen base without casamino acids, 20 g/l dextrose, 5 g/l casamino acids, 100 mM sodium phosphate pH 6.0).

*Growth and expression conditions:* Yeast pre-cultures were performed by inoculating 10 ml of SDCAA medium with one colony from selective agar plate and incubated overnight at 30°C, 200 rpm. The saturated pre-culture (typically $OD_{600}$ of 8–10) was passaged to obtain an initial culture $OD_{600}$ of 0.25-0.50. The culture was grown at 30°C until its $OD_{600}$ reached 0.5-1.0. Cells were centrifuged and re-suspended in 10 ml of SG-CAA galactose induction medium (6.7 g/l yeast nitrogen base without casamino acids, 20 g/l galactose, 5 g/l casamino acids, 100 mM sodium phosphate, pH 6.0) and induced for 16–36 h at 20°C, 200 rpm.

*Yeast labelling and flow cytometry analysis for determining the affinity of CHRD variants for P1.40:*
Cell binding assays were performed according to Hunter *et al.* [18] in order to avoid ligand depletion. $10^5$ cells were taken from the induced culture and washed with 1 ml PBSF (phosphate-buffered saline (PBS), bovine serum albumin (BSA) 0.1%) buffer. Afterwards, cells were resuspended in 0.5 ml of CHRD or CHRD mutants at the desired concentration (0.1 to 200 nM). After incubation at 20°C with shaking (1000 rpm) for 3h, cells were washed with 1 ml ice-cold PBSF to avoid dissociation and resuspended in ice-cold PBSF containing the appropriate fluorescent reporters : 6x-His Epitope Tag antibody Dylight-650 conjugate (Thermo Scientific, catalog number MA1-21315-D650; 1:100 dilution) or HA Tag Monoclonal Antibody APC conjugate (1:50 dilution). Cells were incubated on ice in the dark for 15 minutes and analyzed with BD FACS Aria™ III cytometer. Auto-fluorescence was subtracted for each measure and resulting values were normalized with maximum fluorescence. Data are then fitted with a nonlinear regression analysis to obtain the $K_D$.

**Deep Mutation Scanning (DMS) in CDR3 of P1.40-HA and loss of 6xHis- CHRD binding**

*Library design and generation:* A library of P1.40 variants (region :amino acids 97 to 116) with single amino acid mutations were generated using the "plasmid one pot saturation mutagenesis" method of Wrenbeck *et al.* [19]. After mutagenesis, PCR was performed to recover and amplify the library of P1.40 single mutant genes. The final library was obtained by recombining mutant genes in the YSD plasmid pNT P1.40 between NheI and NotI restriction sites.
*Yeast surface display:* Preparation of competent yeast cells EBY100 (ATCC MYA-4941) and library transformation were performed according to Benatuil *et al* [16] with 2 µg of digested vector and a molar ratio of 25:1 (linear library/digested vector). Gap repair transformations were made in plasmid pNT P1.40 between restriction sites NheI and NotI. Pre-cultures were performed by inoculating 250 ml of SD-CAA medium (6.7 g/l yeast nitrogen base without casamino acids, 20 g/l glucose, 5 g/l casamino acids, 100 mM sodium phosphate, pH 6.0) with 400 µl of transformed cells and incubated overnight at 30˚C, 200 rpm. The saturated pre-culture was passaged in order to obtain an initial culture $OD_{600}$ of 0.25–0.50 in 50 ml. The culture was grown at 30˚C until its $OD_{600}$ reached 0.5–1.0. Cells were centrifuged and re-suspended in 50 ml of SG-CAA galactose induction medium (6.7 g/l yeast nitrogen base without casamino acids, 20 g/l galactose, 5 g/l casamino acids, 100 mM sodium phosphate, pH 6.0) and induced for 16–36 h at 20˚C, 200 rpm.

*Flow cytometry analysis:* For library sorting, $10^7$ induced cells of the library were washed with 1 ml PBSF (phosphate-buffered saline (PBS), bovine serum albumin (BSA) 0.1%) buffer. Cells were resuspended in 2 ml of a solution containing 20 nM CHRD. After incubation at 20°C with shaking (250 rpm) for 3 hours, cells were washed with 1 ml ice-cold PBSF to avoid dissociation. Cells were incubated on ice in the dark for 15 minutes with anti-HA antibody (Invitrogen HA Tag Mouse anti-Tag, DyLight® 650 conjugate, Clone: 2-2.2.14) and anti-6HIS tag antibody (Invitrogen 6x-His Tag Mouse anti-Tag, DyLight® 488 conjugate, Clone: HIS.H8). Cells were subsequently washed with 1 ml ice-cold PBSF and sorted with a BD FACS Aria™ III cytometer using BD FACSdiva™ software. Gates were defined to sort cells with P1.40 expression and marked loss of CHRD binding (Negative sorting). At least 100-fold of the theoretical diversity were sampled, and the gate was collected and recovered for 2 days in SD-CAA medium at 30°C.

*Deep sequencing and analysis of NGS data:* Plasmid DNA of this yeast population was extracted and prepared for sequencing as described in Medina-Cucurella and Whitehead [20]. Two-step PCR was performed to amplify the region of interest and add Illumina adapters and barcodes for multiplexing. Deep sequencing was performed with an Illumina ISeq 100 device (2x150 bp, 300 cycles) with at least 300,000 reads per population. Reads were demultiplexed and the sample was processed separately using the Galaxy platform (https://usegalaxy.org/) using the functions described in Blankenberg *et al*. [21] First, paired reads were joined (Fastq Joiner). A trim was then performed (Fastq Trimmer) on reads to keep just the region of interest in the correct frame. A quality filter (Filter FASTQ) was applied to eliminate reads with a minimum quality score under 30. Next, DNA sequences were translated into protein sequences and identical sequences were grouped. Sequences not repeated at least two times were filtered out. Using the software RStudio, single mutants were selected to allow calculation of enrichment ratios for each single mutation.

**Bio-layer Interferometry (BLI)**

BLI was made with the Pall ForteBio's Octet RED96e System and streptavidin biosensors were hydrated in PBS supplemented with 0.2% BSA and 0.1% Tween20 for at least 10 min before start of each run. Streptavidin biosensors were loaded with different biotinylated proteins. A single run was divided into five distinct steps as follows : (i) Baseline, where the streptavidin biosensor tip was immersed in supplemented PBS for 60 s to obtain a zero baseline; (ii) Loading, where the biotinylated protein was immobilized onto streptavidin-coated biosensor tip for 300 s; (iii) Baseline, where the biosensor tip was again immersed in supplemented PBS for 300 s to remove any unreacted biotinylated protein; (iv) Association, where biotinylated protein was immersed in ligand solutions; (v) Dissociation, where Ligand/ biotinylated protein complex was immersed in supplemented PBS to dissociate the complex. The data from these two last steps was used to estimate differences in interference caused by binding of ligand to biotinylated protein. Blank run was carried out with supplemented PBS and was subtracted from all sample readings. Raw data were pre-processed, analyzed, and fitted by applying the 1:1 binding model as implemented in the manufacturer's Octet Software Software (Data Acquisition 11.0 software and Data Analysis HT 11.0 software, FORTÉBIO).

*PCSK9/P1.40 or CHRD/P1.40:* Binding between PCSK9 (full length) or CHRD with P1.40 was performed with 0.25 µg/ml of biotinylated P1.40. PCSK9 runs were performed with 1600, 800, 400, 200 and 100 nM while CHRD runs were performed with 5, 2.5, 1.2, 0.6 and 0.3 nM. Association time was 3600 s or 1800 s for PCSK9 or CHRD, respectively and dissociation time was 3600 s for both.

*Effects of P1.40 mutations on CHRD interaction:* Streptavidin biosensors were loaded with 1 µg/ml of biotinylated P1.40 variants. CHRD runs were performed with 10, 5, 2.5, 1.2, 0.6 and 0.3 nM for R111A, R43A, E44A, E46A, D62A, D89A and 10 µM for R105A, T109A and F110A. The data from association (300 s) and dissociation (600 s) steps was used to estimate the $K_D$ of CHRD to biotinylated variants P1.40.

*CAP1/PCSK9 or CAP1/CHRD:* Streptavidin biosensors were loaded with 0.2 µg/ml of biotinylated PCSK9 or 0.2 µg/µl of biotinylated CHRD. Runs were performed with 10, 5, 2.5 and 1.2 µM CAP1. The data from association (600 s) and dissociation (600 s) steps was used to estimate the $K_D$ of CAP1 for biotinylated PCSK9 or CHRD.

*Effects of CHRD variants on CAP1 interaction:* Streptavidin biosensors were loaded with 2 µg/ml of biotinylated CAP1. Runs were performed with 10 µM of PCSK9, CHRD or CHRD-2RA or CHRD-3RA. The association and dissociation time were 120 s.

*Effect of an excess of CAP1 on the binding of PCSK9 to P1.40:* Streptavidin biosensors were loaded with 0.25 µg/ml of biotinylated P1.40. The PCSK9/CAP1 complexes was pre-formed overnight at 4°C in PBS with different concentrations of PCSK9 (640, 160 and 80 nM) and 12.3 µM of CAP1.

##### Conformational changes of PCSK9 upon binding of P1.40 assessed by Small-angle X-ray scattering (SAXS)

SAXS experiments were conducted on the SWING beamline at the SOLEIL synchrotron (Gif-sur-Yvette Cedex, FR) [22]. The Supplementary Table 2 resumes the beamline setup and structural SAXS parameters based on the guideline from Trewhella et al [23]. 50 µl of monodisperse PCSK9 protein and complex PCSK9 with P1.40 at a concentration of 2.62 mg/ml and 5.47 mg/ml respectively were injected onto a size-exclusion 5ml column (SEC-3, 300 Å Agilent) using an Agilent HPLC system, and eluted directly into the SAXS flow-through capillary cell at a flow rate of 0.3ml.min$^{-1}$. The elution buffer consisted of 50 mM Tris pH 7.5, 150mM NaCl and 1mM $CaCl_2$. 180 frames were collected during the first minutes of the elution and averaged to account for buffer/substrate scattering for subsequent subtraction from the signal obtained during protein elution. Data reduction to absolute units, frame averaging, and subtraction were done using FOXTROT (Synchrotron SOLEIL) and US-SOMO software [24]. The frames with a stable Rg under the main pics were averaged to obtain the sample SAXS curve. Data were then analysed using the ATSAS 3.0.1 software suite [14,25]. Three starting atomic models were generated using MODELLER [26] and PDB structures 2P4E and 7ANQ as templates. The first two PCSK9 models were a closed conformation with the CHRD domain interacting with the catalytic domain and an extended conformation no interaction between these domains. The extended conformation was then used as a model for the construction of the third model corresponding to the PCSK9/P1.40 complex. DADIMODO software [10] was used to refine these three models according to their SAXS curves. Fifteen models were generated and ranked according to their $\chi 2$ for each DADIMODO calculation (**Supplementary Table 2**).

##### Immunoprecipitation

CAP1 (HA), PCSK9 and PCSK9-Δ33-58 conditioned media were produced from HepG2-*PCSK9*$^{-/-}$ cells transfection of corresponding cDNA constructs. 24 h post-transfection, the cells were incubated for 1 h with SFM followed by 18 h incubation in SFM containing or not 1 µg/ml of P1.40 nanobody. The proteins were then immunoprecipitated from 2 ml of media after 2 h incubation with HA-coupled to agarose beads (Santa Cruz). The beads were washed twice with lysis buffer, heated to 95 °C in SDS sample buffer, resolved by SDS-PAGE, transferred to nitrocellulose membranes for Western blotting and analyzed as described above.

##### Cycloheximide chase assay

Briefly, HepG2 PCSK9$^{-/-}$ cells were subjected to siRNA silencing as described above then incubated with 40 µg/ml cycloheximide (Sigma) for 18h as described previously [27]. Cell samples collected at time points ranging from 0 to 6 h were resolved and analysed by WB.

##### Inhibition of protein expression by small-interfering RNAs (siRNAs)

siRNA analysis was performed using INTERFERin® (PolyPlus) transfection reagent according to manufacturer's instruction. siRNAs (50 nM final concentration) targeted against: CTL [non targeting sequence, siGENOME Non-Targeting siRNA Pool #1, Catalog#: D-001206-13-05]; CAP1 [siGENOME Human CAP1 (10487) siRNA – SMARTpool, Catalog#: M-012210-01-0005]; CHC [ON-TARGETplus Human CLTC (1213) siRNA – SMARTpool, Catalog#: L-004001-01-0005]; and CAV1 [ON-TARGETplus Human CAV1 (857) siRNA – SMARTpool, L-003467-00005] were purchased from Dharmacon (Horizon Discovery). Gene silencing efficiency was assessed by Western blotting.

##### Statistical analysis

Statistics were performed with GraphPad Prism (9.4.1) using appropriate statistic tests (i.e., Welch's *t*-test or one-way/two-way ANOVA, coupled to Tukey's multiple comparisons test) to assess the significance of all data sets as previously described [28]. Data are presented as means +/- SEM and a $p$ values lower than 0.05 were considered as statistically significant.

### Supplementary Figures

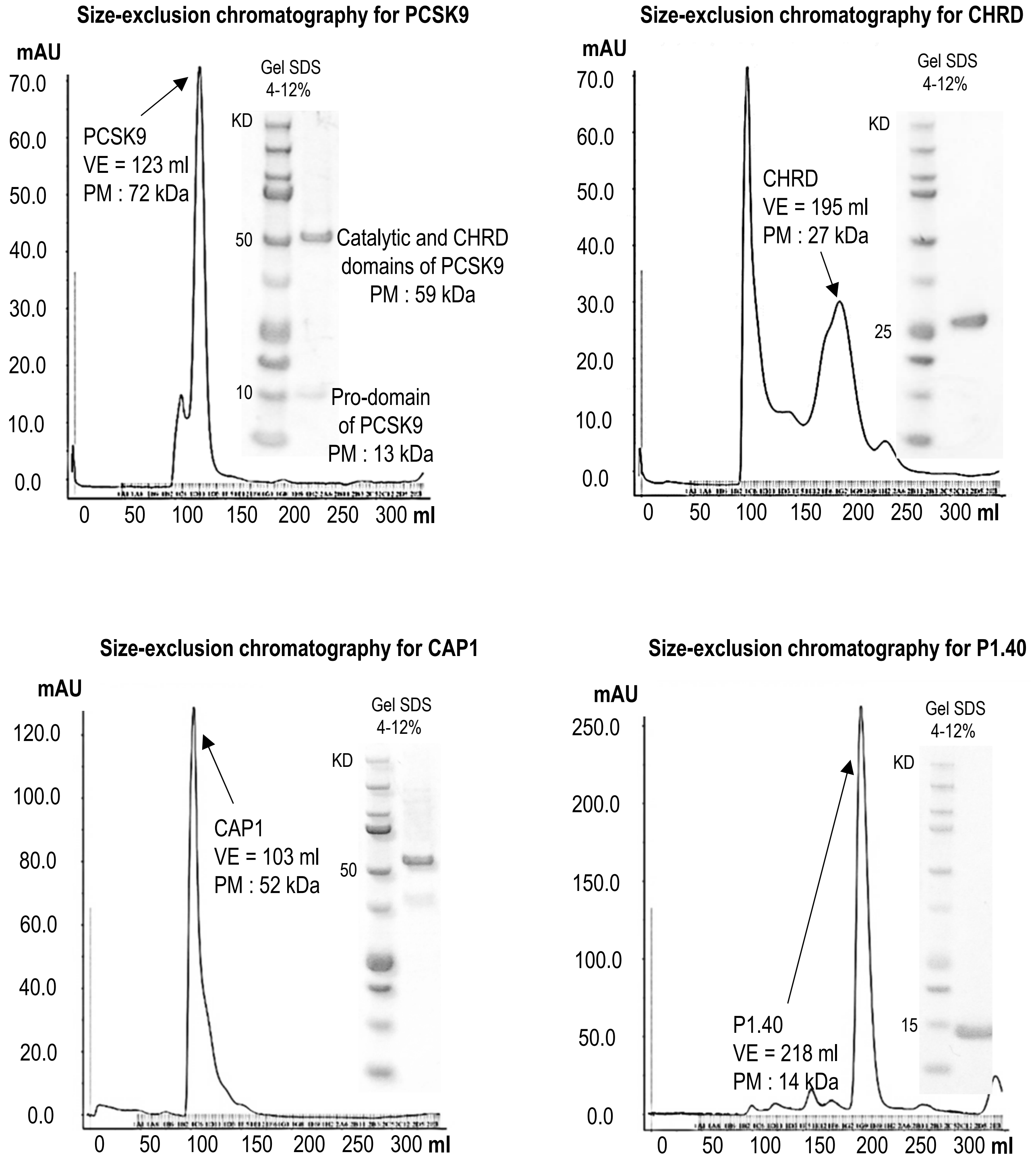


**Supplementary Figure 1** : PCSK9, CHRD, CAP1 and nb P1.40 purified from Size-exclusion chromatography (SEC) Sephacryl ® S-100 HR GE Healthcare in buffer 50 mM Tris HCl pH 8, 150 mM NaCl. and loaded on 4-12% SDS PAGE.

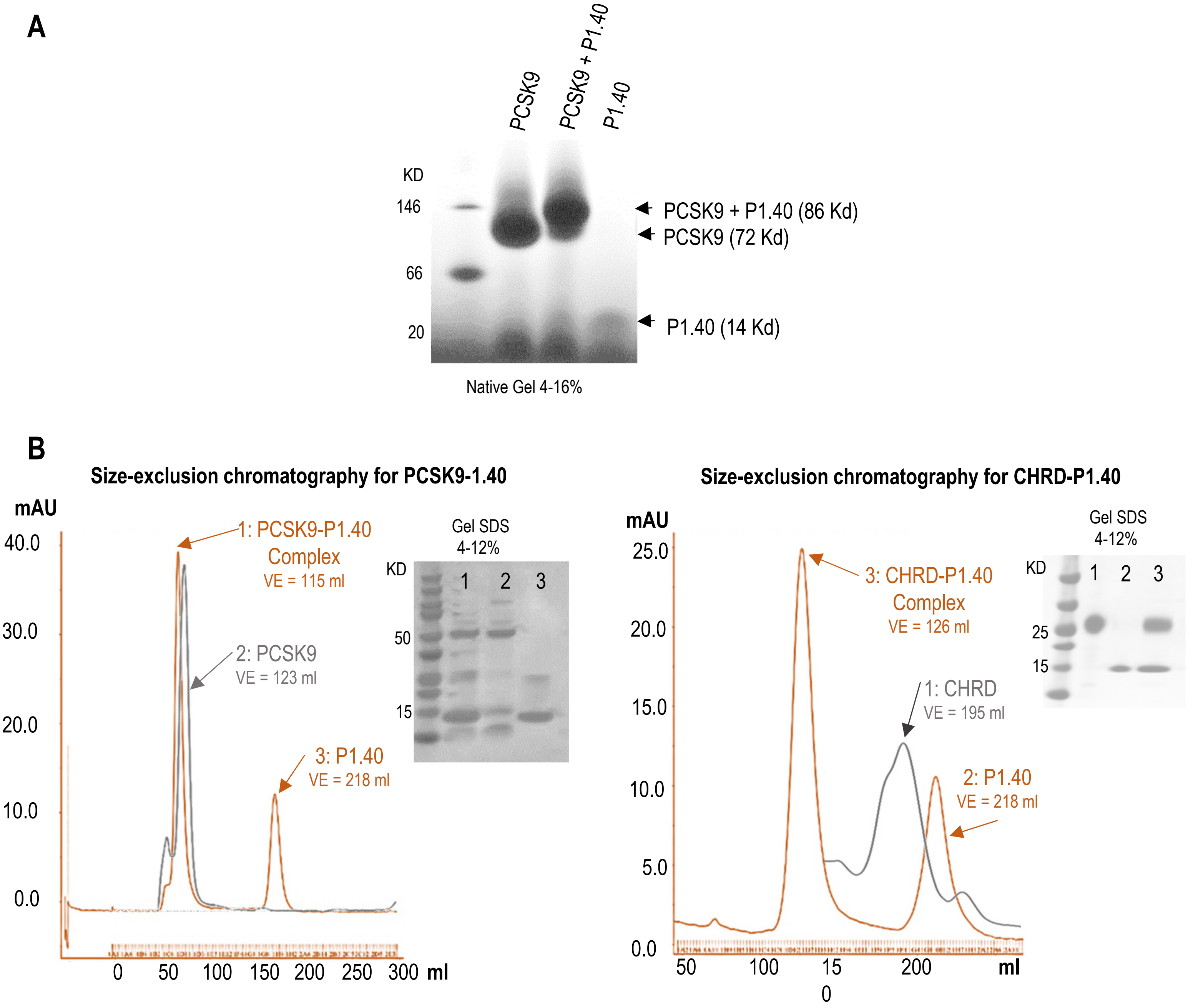


**Supplementary Figure 2**: **Characterization of the PCSK9-P1.40 complex. (A)** NativePage$^{TM}$ 4-16% Bis-Tris Ge. (**B**) Purification of PCSK9-P1.40 and CHRD-P1.40 by Size-exclusion chromatography (SEC) Sephacryl ® S-100 HR GE Healthcare in buffer 50 mM Tris HCl pH 8, 150 mM NaCl and loaded on 4-12% SDS PAGE.

**A**

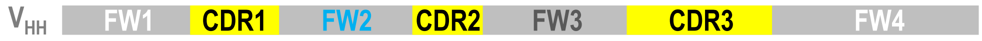


**B**

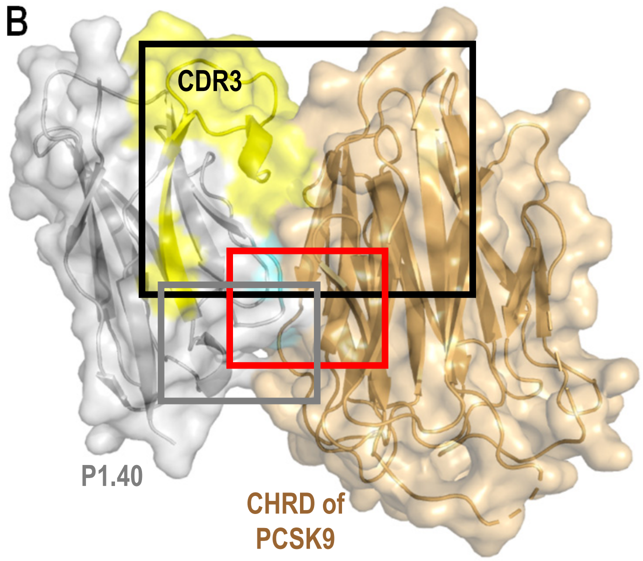


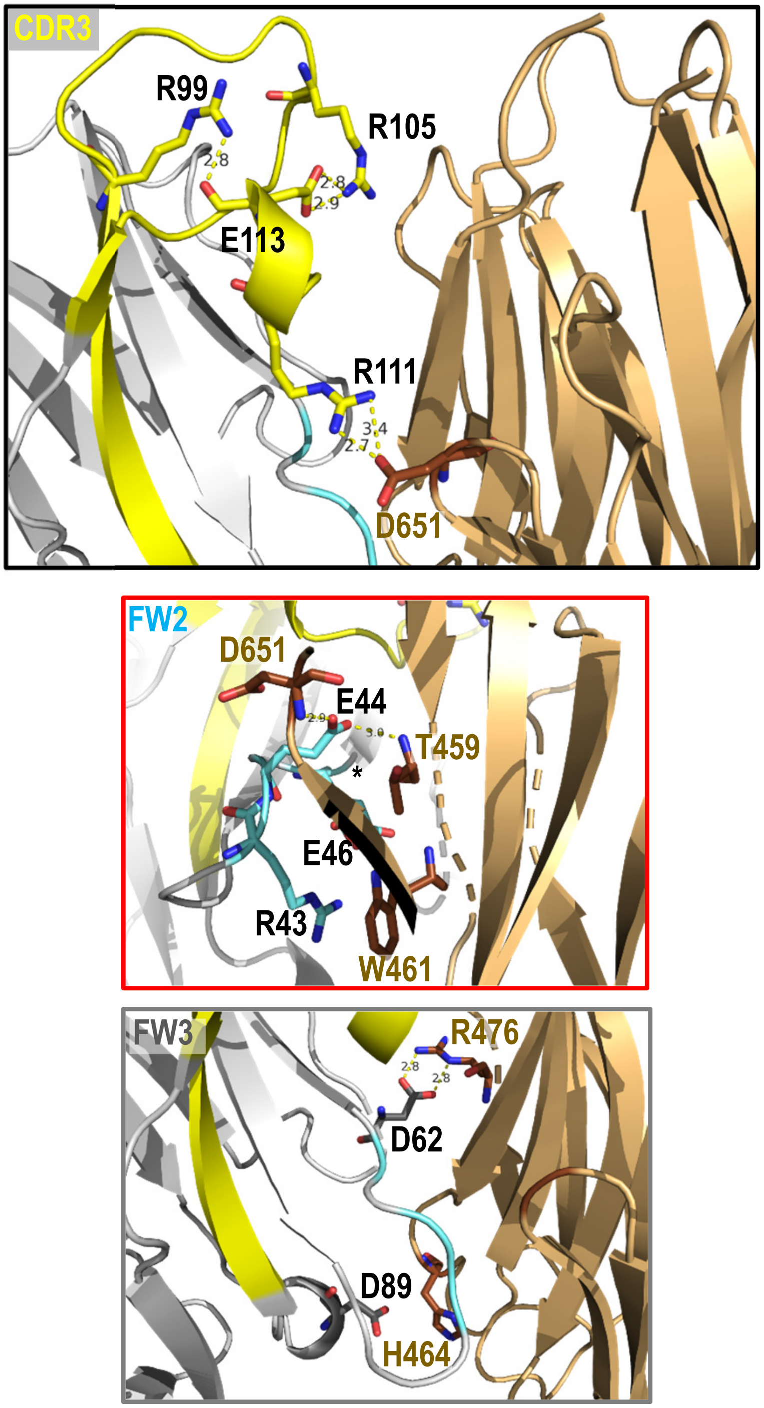


| Minor interactions | | | |
|---|---|---|---|
| **CDR3** | **M3** | **Interaction** | **Distance (Å)** |
| R111 | D651 | H-bond | $NH_2^{\eta}$---$O^{\delta}$ 2.78 |
| | | Polar | $NH_2^{\eta}$---$O^{\delta}$ 3.4 |
| **FW2** | **M1** | **Interaction** | **Distance (Å)** |
| R43 | W461 | Polar | $NH_2^{\eta}$ ---NH |
| E44 | T459 | H-bond | $O^{\varepsilon}$---$NH_2$ 3.0 |
| E46 | T459 | Polar | $O^{\varepsilon}$---O 3.9 |
| E46 | W461 | Polar | $O^{\varepsilon}$---NH 3.5 |
| | | | $O^{\varepsilon}$---NH 4.0 |
| **FW2** | **M3** | **Interaction** | **Distance (Å)** |
| E44 | D651 | H-bond | $O^{\varepsilon}$---$NH_2$ 2.9 |
| **FW3** | **M1** | **Interaction** | **Distance (Å)** |
| D62 | R476 | H-bonds | $O^{\delta 1}$---$NH_2^{\eta}$ 2.8 |
| | | | $O^{\delta 2}$---$NH^{\varepsilon}$ 2.8 |
| D89 | H464 | Polar | $O^{\delta}$---$NH^{\delta}$ 5.1 |

**Supplementary Figure 3: Analysis of P1.40-CHRD interactions in the crystal complex PDB 7ANQ**. (**A**) Structure of the variable heavy chain-only ($V_{HH}$) antibody P1.40 raised in llama [8]. The antigen binding site is constituted of three complementary-determining regions (CDRs) framed with framework regions (FWs) implicated in the core structure of the antibody. (**B**) Surface representation of the P1.40 (grey)-CHRD (light brown) crystal complex and modeling of additional interactions established between P1.40 domains CDR3 (yellow), FW2 (cyan) and FW3 (dark grey) and the CHRD of PCSK9, which is composed of M1, M2 and M3 subdomains. **CDR3** (aa 105-122): between $Arg_{111}$ and M3 $Asp_{651}$. **FW2**: between $Arg_{43}$, $Glu_{44}$, $Glu_{46}$ and M1 $Thr_{459}$, $Trp_{461}$ or M3 $Asp_{651}$. **FW3**: between $Asp_{62}$, $Asp_{89}$ and M1 $His_{464}$, $Arg_{476}$.

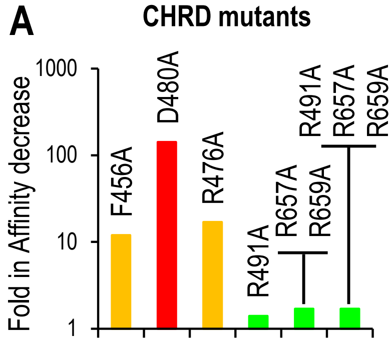


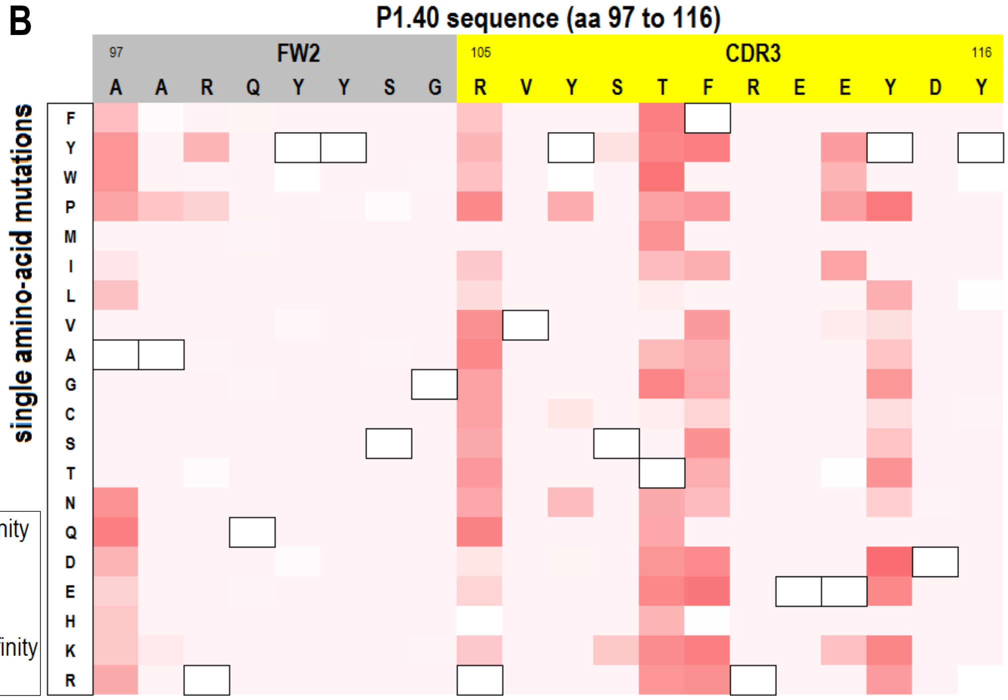


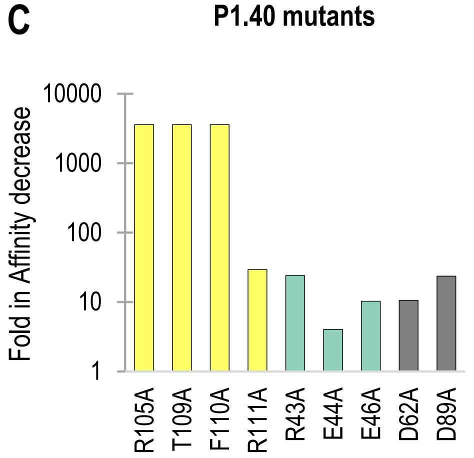


| P1.40 mutation | ka (1/Ms) | ka Error | kdis (1/s) | kdis Error | KD (M) | Fold affinity decrease |
|---|---|---|---|---|---|---|
| **WT** | 3.76E+05 | 2.95E+02 | 3.49E-05 | 1.22E-07 | 9.03E-11 | |
| **R105A** | > 1.00E+03 | - | >1.00E-3 | | > 1.00E-06 | >10000 |
| **T109A** | > 1.00E+03 | - | >1.00E-3 | | > 1.00E-06 | >10000 |
| **F110A** | > 1.00E+03 | - | >1,00E-3 | | > 1.00E-06 | >10000 |
| **R111A** | 4.20E+05 | 3.18E+03 | 1,47E-03 | 2.21E-06 | 3.49E-09 | 38.7 |
| **R43A** | 4.95E+05 | 1.91E+03 | 1,41E-03 | 2.62E-06 | 2.86E-09 | 31.7 |
| **E44A** | 7.53E+05 | 1.66E+03 | 3,59E-04 | 1.23E-06 | 4.77E-10 | 5.3 |
| **E46A** | 5.26E+05 | 3.13E+03 | 6,40E-04 | 3.66E-06 | 1.22E-09 | 13.5 |
| **D62A** | 5.30E+05 | 1.84E+03 | 6,68E-04 | 2.31E-06 | 1.26E-09 | 13.9 |
| **D89A** | 3.16E+05 | 4.94E+02 | 8,82E-04 | 9.80E-07 | 2.79E-09 | 30.95 |

**Supplementary Figure 4: Identification of the key interacting residues in the P1.40-CHRD complex.** (**A**) Decreased affinity was measured by yeast cell-surface display of P1.40 and CHRD binding after Ala replacement of selected CHRD residues. (**B**) A library containing every possible amino acid change at each position of a selected region of P1.40 (Deep Mutational Scanning) was generated by PCR and tested for the loss of capacity of each P1.40 mutant expressed at the yeast cell surface to bind the CHRD (negative sorting). White color indicates that P1.40-CHRD binding was unchanged while the darkest red indicate a loss of binding. (**C**) Impact of targeted mutations on biotinylated P1.40 affinity for the CHRD are shown. Corresponding kinetic constants were analyzed by BLI using a streptavidin sensor.

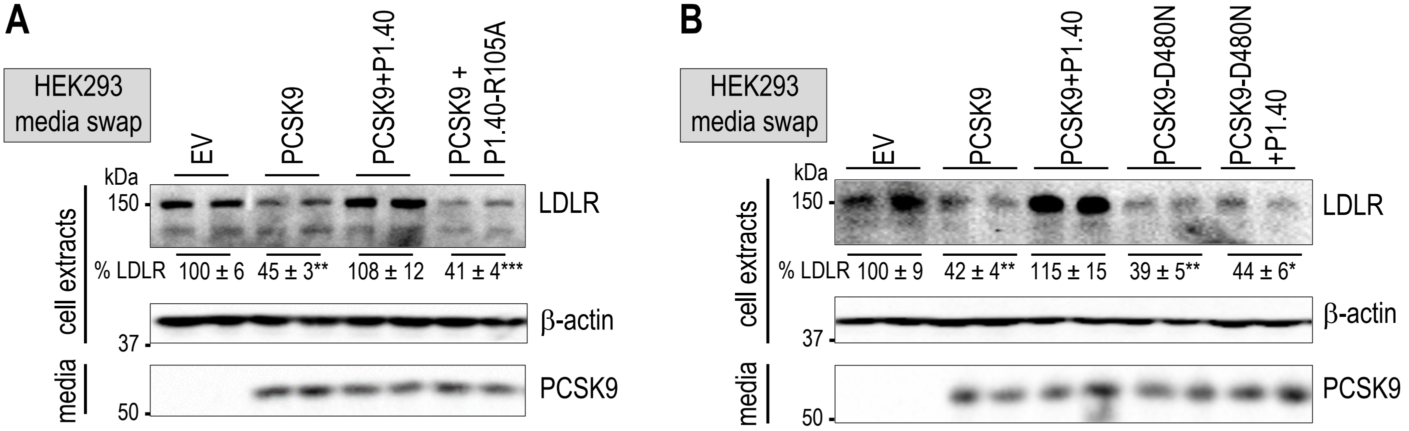


**Supplementary Figure 5: Cell validation of the key interaction between the $Arg_{105}$ in P1.40 and $Asp_{480}$ for the inhibitory effect of P1.40.** (**A**) HepG2-*PCSK9$^{-/-}$*cells were incubated with conditioned media from HEK293 cells expressing an empty vector (EV) or PCSK9-V5 (0.3 µg/ml), in the absence or presence of purified P1.40 or P1.40-R105A mutant (1 µg/ml). LDLR protein levels were normalized to β-actin and set to 100 for EV. (**B**) The same conditions as in (A) were used, except that PCSK9 or PCSK9-D480N were expressed and used at 0.3 µg/ml. Means ± SEM for three (A) or two (B) independent experiments. *P* values for LDLR (*, *P*<0.05; **, *P*<0.01; ***, *P*<0.001) were obtained by two-way ANOVA coupled to Tukey's multiple comparisons tests.

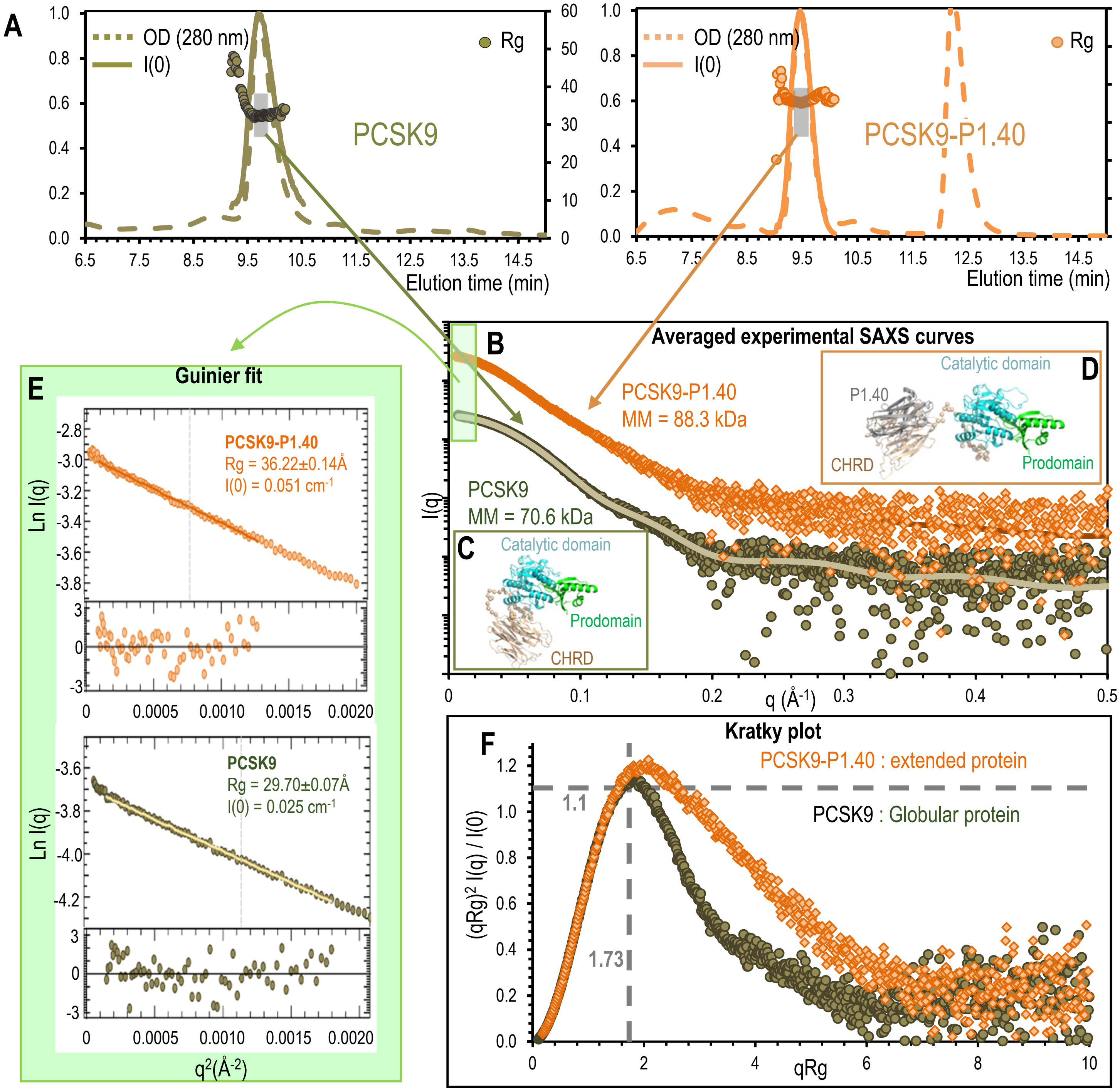


**Supplementary Figure 6:** Size exclusion chromatography and SAXS data. (**A**) UV (280 nm), I(0) (vector intensity; solid) and Rg (radius of gyration; dots) elution profiles of the protein PCSK9 alone (green) and in complex with P1.40 (orange). Normalized UV and I(0) values are shown on the left y-axis and Rg values in Å on the right one. $OD_{280nm}$ profiles indicates that PCSK9 is monomeric in solution, peaking at 9.8 min, and that the PCSK9/P1.40 complex is stable, does not dissociate and peaks at 9.5 min. The second peak at 12.3 min corresponds to excess monomeric P1.40. Selected fractions for SAXS curves averaging (9.5 to 10 min and 9.3 to 9.7 min for PCSK9 alone and in complex, respectively) are shown in grey. (**B**) Experimental SAXS curves were obtained by averaging and defined scattering vectors in $Å^{-1}$ (x-axis) and their intensities (y-axis). The CRYSOL [9] results fitting (green and orange lines) with the best DADIMODO [10] models for PCSK9 alone (**C**) and in complex with P1.40 (**D**) gave a $\chi^2$ of 1.15 and 0.97 respectively. (**E**) The light green rectangle in (B) corresponds to the selected area for the Guinier fit [11] for q closed to 0. The Guinier law defines the radius of gyration Rg and I(0). (**F**) In the normalized Kratky plot [12,13] making it possible to quickly assess the globular nature and the flexibility, the dashed cross represents the theoretical value for the maxima of a globular protein (peaks at 1.1 for qRg of 1.73). The experimental curves (dot) show that the global shape of the complex with a maximum peak at 1.2 (orange symbols) is more extended compared to the global shape of the protein PCSK9 alone (green symbols) with a maximum at 1.16. The estimation of molecular weight of PCSK9 alone (70.6 kDa) and of the PCSK9/P1.40 complex (88.3 kDa) by SAXS using the Bayesian estimation from ATLAS [14] indicates that a single protein P1.40 (14.4 kDa) binds to PCSK9 (72.5 kDa) in solution.

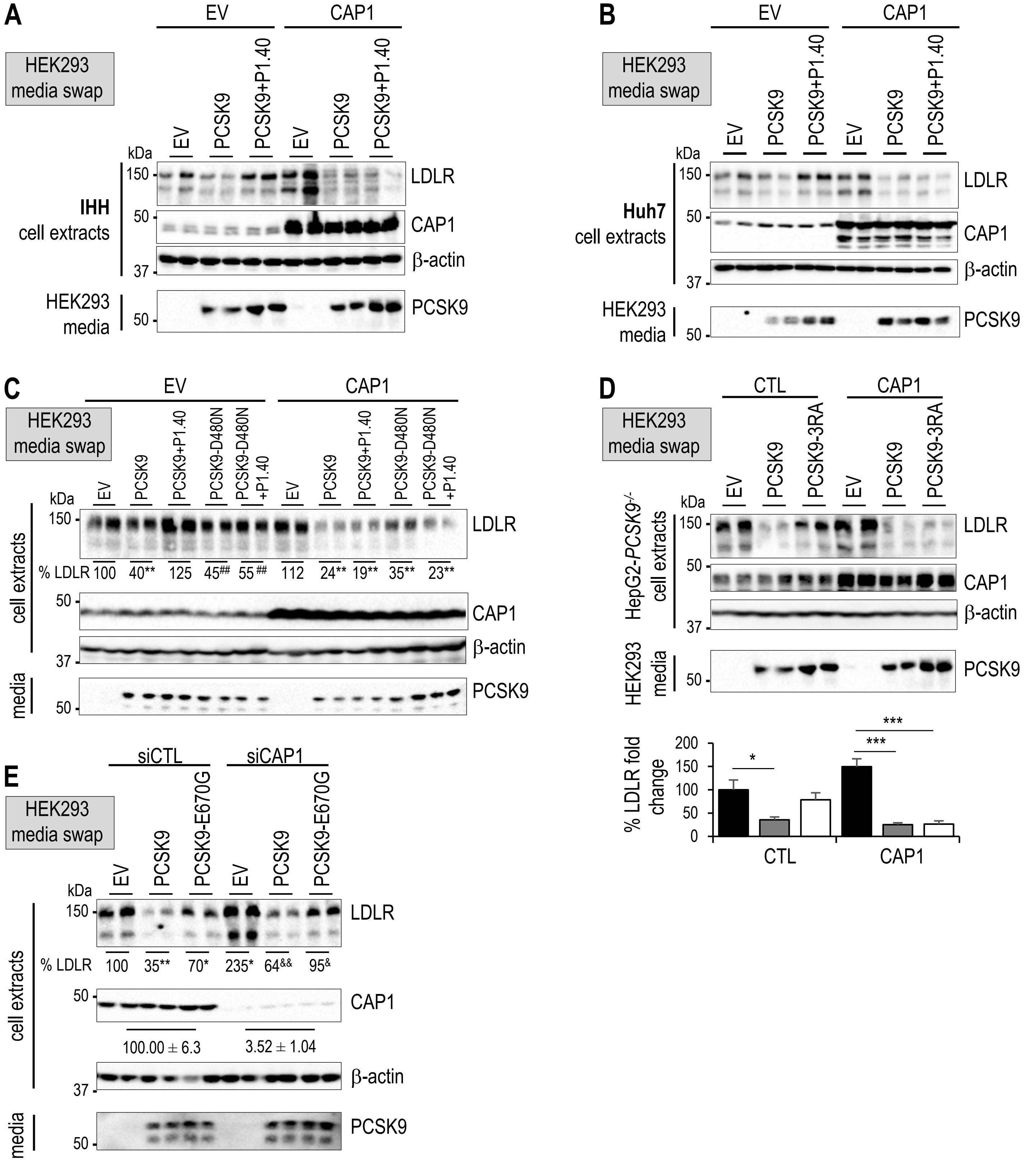


**Supplementary Figure 7: Regulation of PCSK9 activity by CAP1 and P1.40.** IHH (**A**) or Huh7 (**B**) cells were transfected for 24 h with an empty- (EV) or CAP1-expressing vector and then exposed for 18 h to conditioned media from HK293 cells, as in Figure 5A. (**C**) HepG2-*PCSK9*$^{-/-}$ cells expressing an empty vector (EV) or CAP1 (24 h) were treated for 18 h with conditioned media from HEK293 cells expressing an empty vector (EV) or incubated with PCSK9 or PCSK9-D480N (0.3 µg/mL) in combination with P1.40 (1 µg/ml). (**D**) HepG2-*PCSK9*$^{-/-}$ cells were treated under the same conditions as in (C) with PCSK9 or PCSK9-3RA. (**E**) HepG2-*PCSK9*$^{-/-}$ cells were transfected with siRNA constructs including siCTL or siCAP1 and incubated with conditioned media from HEK293 cells expressing an empty vector (EV), PCSK9 or PCSK9-E670G mutant (0.3 µg/ml) for 18 h. Cell extracts and media were analyzed by Western blot. Representative blots of three (A-D) and two (E) independent experiments are shown. Data represent means ± SEM (*, $P$<0.05; ***, $P$<0.001; $^{\&}$, $P$<0.05; $^{\&\&}$, $P$<0.01) obtained by two-way ANOVA coupled to Tukey's multiple comparisons tests.
